# Colibactin produced by a honeybee symbiont defends against pathogens and shapes the gut community

**DOI:** 10.64898/2025.12.27.695564

**Authors:** Yulin Song, J. Elijah Powell, Joel W. H. Wong, Yunxi Liu, Miguel A. Aguilar Ramos, Tyler De Jong, Patrick J. Lariviere, Jayaditya Maganti, Emily P. Balskus, Nancy A. Moran

**Affiliations:** Department of Integrative Biology, University of Texas, Austin, TX 78712, USA; Department of Chemistry and Chemical Biology, Harvard University, Cambridge, MA 02138, USA; Department of Molecular Biosciences, University of Texas, Austin, TX 78712, USA; Howard Hughes Medical Institute, Harvard University, Cambridge, MA 02138, USA

**Keywords:** Colibactin, gut microbiota, interbacterial competition, honeybee, DNA damage

## Abstract

Colibactin is a bacterial genotoxin that is linked to colorectal cancer and implicated in interbacterial competition; however, its role in natural communities remains unknown. *Frischella perrara*, a symbiont living only in honeybee guts, produces colibactin and causes DNA damage. Here, we found that *F. perrara* mono-colonization reduces bee lifespan but increases survivorship following challenge with an opportunistic pathogen. *F. perrara* reduces pathogen loads by causing colibactin-dependent DNA damage and prophage induction. *clbS*, a gene conferring protection from colibactin, is ubiquitous among bacteria restricted to bee guts but is absent from opportunistic colonizers including bee pathogens. Our findings provide evidence that colibactin functions in interbacterial competition, exerts selective pressure on community members, and confers protection against pathogens, while permitting co-evolved gut symbionts to persist.

## Introduction

The intestinal tracts of animals provide distinctive habitats for microbial communities. Host-associated microorganisms establish structured consortia, as found in the human gut, where they may benefit hosts via multiple routes, including nutrient metabolism, immunomodulation, and pathogen defense (*1*). Besides influences from host factors, the commensal gut microbiota is shaped by intricate interbacterial interactions (*2*). Antagonistic interactions among bacterial strains or species within the community can be mediated by contact-dependent weapons or secreted antimicrobial compounds, which have central roles in competition for nutrients and niches (*3, 4*). Such compounds have the potential to deter invading pathogens, thus affecting host health.

The western honeybee, *Apis mellifera*, serves as the most important pollinator in agriculture and has recently been developed as a model for gut microbiota research (*5*). It has a relatively simple gut microbiota which is dominated by five core bacterial taxa: *Snodgrassella*, *Gilliamella*, *Bombilactobacillus*, *Lactobacillus*, and *Bifidobacterium* (*6*). These taxa are spatially organized in the hindgut (*6, 7*), with considerable strain-level divergence in natural communities (*8*). Other non-core bacterial symbionts, such as *Bartonella* spp. and *Frischella perrara*, are restricted to bee guts and common in honeybee populations but are not ubiquitous. *F. perrara* is present in almost all hives and in approximately half of the bees (24–82%) (*9*). Some environmental, opportunistic pathogens, such as *Serratia marcescens* (*10*), are present in low abundance in both the bee gut and the hive environment (*5*).

*F. perrara* densely colonizes the pylorus, a short region between the midgut and hindgut, overlapping with a bacterial biofilm formed by *Snodgrassella alvi*, *Gilliamella apicola* and *Gilliamella apis* in the ileum (*6, 9*). Vigorous colonization by *F. perrara* activates bee immune responses (*11*) and causes a melanized scab-like phenotype in this region (*12*). Intriguingly, *F. perrara* possesses a colibactin genomic island (denoted *pks* or *clb*) for which the genetic organization is conserved with other bacteria possessing this pathway (*12*)(Fig. 1A), including certain *Escherichia coli* and other *Enterobacteriaceae* strains present in the human gut (*13, 14*), *Pseudovibrio* FO-BEG1 isolated from a marine coral (*15*), and *Erwinia oleae* from tree knots (*16*). The colibactin locus is likely a part of the core genome of *F. perrara* (see Supplementary Text and Supplementary Fig. S1). The scab-like phenotype is not caused by colibactin, as a gene deletion mutant, *ΔclbB*, that cannot produce colibactin still triggers this phenotype (*17*).

**Fig. 1.**
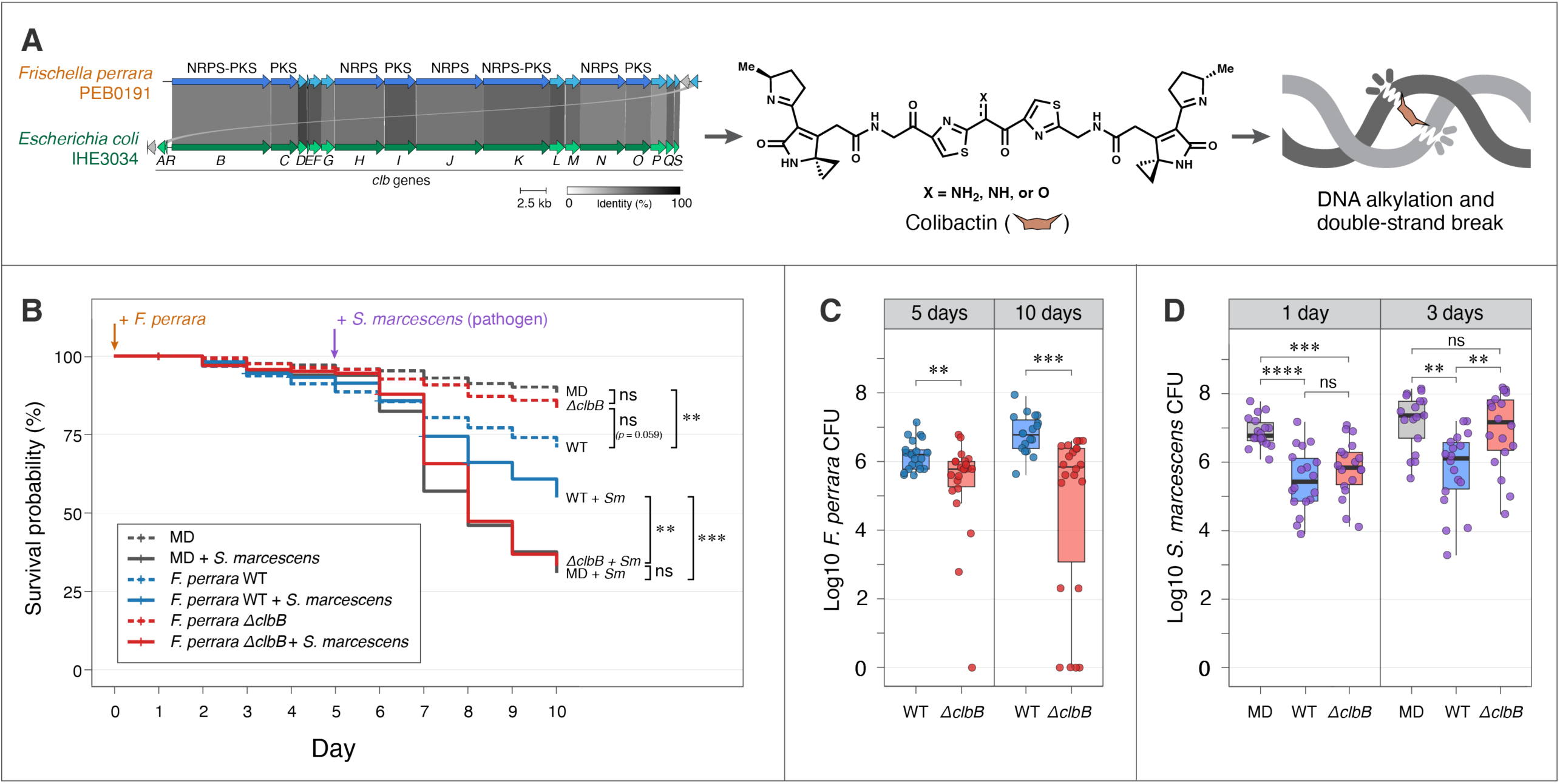
The honeybee gut symbiont *Frischella perrara* can be beneficial or detrimental to hosts, and outcomes depend on colibactin production and pathogen presence. (A) *F. perrara* encodes a genomic island (*pks* or *clb*) which has conserved synteny and high sequence homology to that of some *Escherichia coli* strains from human guts and produces the DNA-damaging agent colibactin. Genes encode nonribosomal peptide synthetase (NRPS), polyketide synthase (PKS), or NRPS-PKS hybrid enzyme of the colibactin biosynthetic pipeline are in dark blue or green, and other accessory genes are in light blue or green. Homologous genes are linked with connections, the greyscale of which reflects percent of amino acid identity as indicated by the color key. Gene names (*clbA–S*) are given below the synteny. The *F. perrara* genes/proteins are detailed in the GenBank annotation (accession: NZ_CP009056.1). (B) Survival of gnotobiotic bees inoculated with *F. perrara* WT, *ΔclbB* (colibactin null mutant) or not inoculated (microbiota-deprived control, MD), and challenged with a pathogen *Serratia marcescens* or not after 5 days. The Kaplan-Meier survival curves combined three independent experiments with bees sourced from different hives (∼20 bees per cup cage, 3 cages per group in each experiment). ANOVA test (*χ^2^* = 185.38, *df* = 5, *p* < 0.0001, *n* = 1080) showed statistically significant difference among the treatment groups. Post-hoc pairwise comparisons were conducted using estimated marginal means. (C) CFUs showing the abundance of *F. perrara* WT and *ΔclbB* in the whole gut 5 and 10 days after inoculation (total *n* = 83). Statistical comparisons were conducted using Wilcoxon rank sum test. (D) CFUs showing the abundance of *S. marcescens* in the abdomen of bees 1 day and 3 days after challenging. The bees were pre-inoculated with *F. perrara* WT, *ΔclbB* or not inoculated. Kruskal-Wallis test showed statistical significance among the groups in both day 1 (*χ^2^* = 23.426, *df* = 2, *p* < 0.0001, *n* = 54) and day 3 (*χ^2^* = 14.425, *df* = 2, *p* < 0.001, *n* = 53). Post-hoc pairwise comparisons were conducted using Dunn’s test. Statistical significance: ****, *p* < 0.0001; ***, *p* < 0.001; **, *p* < 0.01; *, *p* < 0.05; ns, not significant.

Colibactin is a hybrid nonribosomal peptide synthetase-polyketide synthase (NRPS-PKS) product which causes double-strand breaks in DNA via alkylation and interstrand crosslink formation (*13, 18–21*), leading to cell cycle arrest in eukaryotic cells and tumor formation (*13, 22–26*). Like *E. coli*, *F. perrara* imposes *clb*-dependent genotoxic damage on eukaryotic cells (*12*). Further, in vitro and in vivo studies show that colibactin production by *E. coli* damages DNA and inhibits the growth of other bacterial species (*27–31*), influences community structure (*28, 32*), and may enhance colonization in the gut (*33*). However, few studies have addressed the role of colibactin within natural gut communities.

Honeybees offer the opportunity to use newly emerged, microbiota-deprived adult bees, with similar age and genetic background, for defined inoculations and further tests (*34*). In this study, we investigated the effects of *F. perrara* colonization and colibactin production on other bacterial community members and on bee health.

## Results

### *F. perrara* has dual effects on honeybee health

To assess the influence of *F. perrara* colonization on honeybee health, we carried out a pilot experiment. Gnotobiotic bees were pre-inoculated with *F. perrara* and, after 5 days, challenged with an opportunistic pathogen, *S. marcescens* N10A28, a strain originally isolated from a honeybee gut that can lower bee survivorship (*35*). Bees mono-inoculated with *F. perrara* did not differ significantly in mortality compared to microbiota-deprived (MD) controls (*p* = 0.072) (Supplementary Fig. S2). When challenged with the pathogen, however, *F. perrara*-inoculated bees had much higher survivorship than MD bees (*p* = 0.0068).

We wondered if colibactin production underlies *F. perrara*’s protective effect. In a further experiment, we included strain *F. perrara ΔclbB*, which has a deletion in a NRPS-PKS gene that is essential for colibactin biosynthesis (*17*). We tested bees sourced from different hives and inoculated with *F. perrara* WT, *F. perrara ΔclbB* or not inoculated, and subsequently challenged with *S. marcescens* or not challenged. Bees inoculated with *F. perrara* WT died sooner than MD bees in the absence of pathogen (*p* = 0.0014), and, as in the previous experiment, inoculation with *F. perrara* WT improved survivorship after pathogen challenge (*p* = 0.0002) (Fig. 1B). Mono-inoculation with *F. perrara ΔclbB* had no significant effect on survival compared to MD (*p* = 0.82). *F. perrara ΔclbB* failed to protect against *S. marcescens* (*p* = 0.997), whereas WT provided protection against the pathogen (*p* = 0.0016), as in the earlier experiment. For the mono-inoculated bees over a longer time frame (18 days), mortality of bees with *F. perrara ΔclbB* was higher than that of MD bees (*p* = 0.025) but lower than for WT-inoculated bees (*p* = 0.014) (Supplementary Fig. S3).

We plated samples and counted colony-forming units (CFUs) to estimate abundances of *F. perrara* or *S. marcescens* in bee guts. In gnotobiotic bees, *F. perrara* WT showed better colonization than *ΔclbB* after 5 and 10 days (Fig. 1C), consistent with previous observations (*17*). Pre-inoculation with either *F. perrara* WT or *ΔclbB* reduced gut levels of *S. marcescens* 1 day after challenge. This colibactin-independent short-term protection most likely results from symbiont-mediated stimulation of host immune responses including reactive oxygen species (ROS; see Supplementary Text and Supplementary Fig. S4). However, after 3 days, only WT-inoculated bees had reduced pathogen loads while *ΔclbB*-inoculated bees had loads similar to those of MD bees (Fig. 1D), linking long-term pathogen suppression to colibactin production. Taken together, these results suggest that colibactin specifically confers host protection against the pathogen.

### *F. perrara* suppresses pathogens via colibactin-dependent DNA damage

Since *F. perrara* reduced *S. marcescens* proliferation in a colibactin-dependent way, we investigated possible mechanisms for this inhibition. To investigate bacterial DNA damage caused by colibactin, a cassette with eGFP under the promoter region of the *S. marcescens recA* gene was inserted onto the plasmid pSL1-E2C (E2C confers crimson fluorescence), generating a new plasmid pSL1-E2C-N10P*_recA_*eGFP for DNA damage reporting (Fig. 2A). *S. marcescens* carrying this plasmid gave constitutive crimson fluorescence and inducible green fluorescence, triggered by DNA damage after treatment with the mutagen mitomycin C (MMC) (Supplementary Fig. S5). A flow cytometry assay elaborated that the GFP signal induced by MMC was concentration-dependent (Supplementary Fig. S6, S7).

**Fig. 2.**
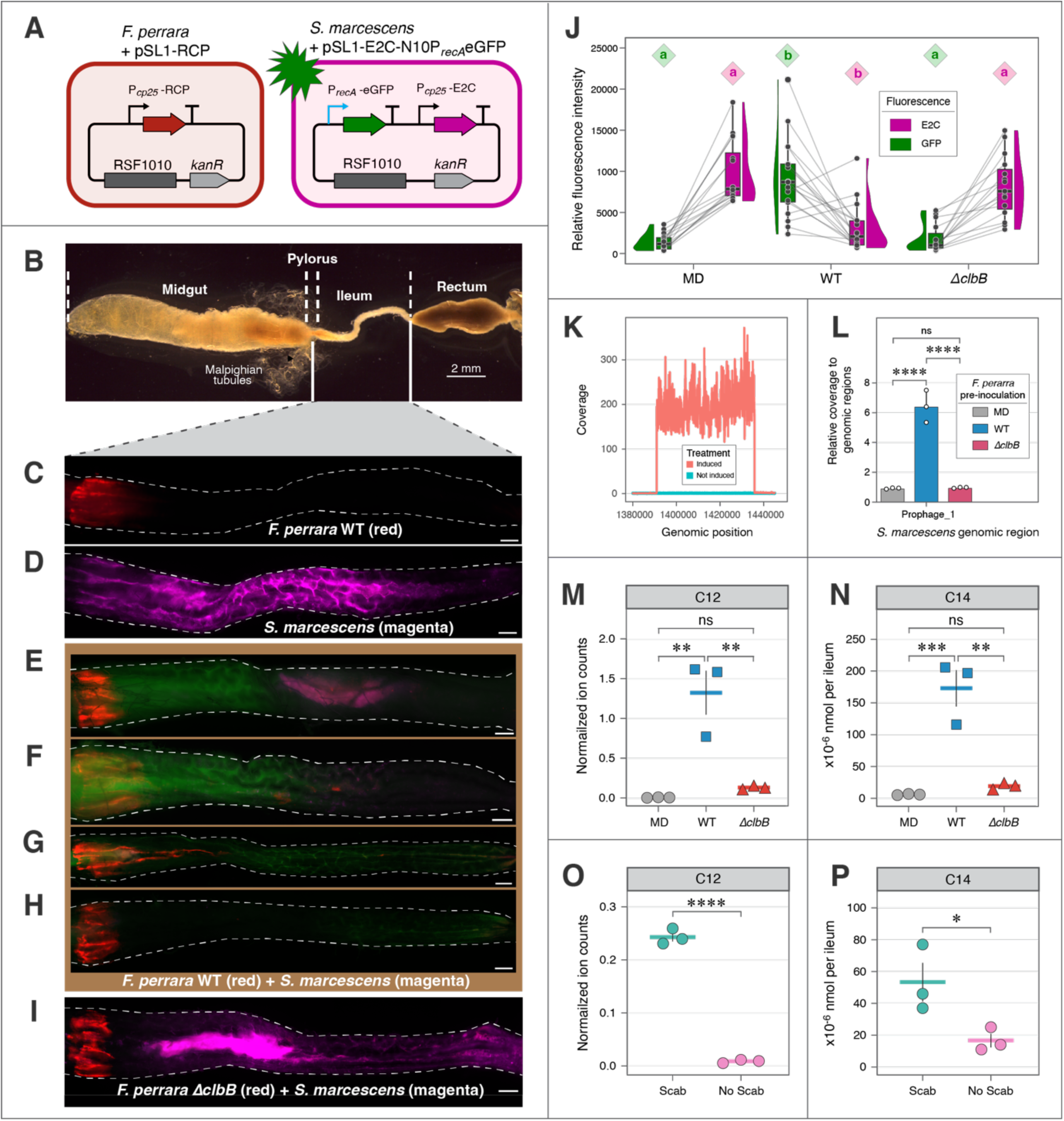
Colibactin production by *F. perrara* suppresses *S. marcescens* pathogen in honeybee hindguts by causing DNA damage and prophage induction. (A) Schematic representation of strains with reporter plasmids. *F. perrara* WT or *ΔclbB* carrying pSL1-RCP that constitutively expresses a red chrome protein. *S. marcescens* carrying pSL1-E2C-P*_recA_*eGFP that expresses E2-Crimson constitutively and GFP inducibly, triggered by DNA damage. (B) Gut structure of a dissected adult honeybee. (The image shows two stitched photos.) (C) Ileum mono-colonized with *F. perrara* WT. (D) Ileum mono-colonized with *S. marcescens*. (E–H) Ilea pre-colonized with *F. perrara* WT and subsequently challenged with *S. marcescens*. (I) Ileum pre-colonized with *F. perrara ΔclbB* and challenged with *S. marcescens*. Scale bars = 0.2 mm in C–I. *F. perrara* was fed to newly emerged microbiota-deprived bees at day 0, and *S. alvi* was fed at day 6; fluorescence was checked after another two days. (J) Measured mean fluorescence intensities in the ilea of gnotobiotic bees pre-colonized with *F. perrara* WT, *ΔclbB* or not inoculated (MD) and challenged with *S. marcescens*. Each pair of connected dots shows intensity of GFP (DNA damage) and E2C (*S. marcescens*) in an individual bee. Kruskal-Wallis test indicated statistical significance among treatment groups in GFP intensity (*χ^2^*= 30.1, *df* = 2, *p* < 0.0001, *n* = 49) and E2C intensity (*χ^2^* = 23.3, *df* = 2, *p* < 0.0001, *n* = 49). Post-hoc pairwise comparisons were conducted using Dunn’s test. Groups with different letters are significantly different (*α* = 0.05). (K) Change in depth of sequencing reads for a prophage-like region (genomic coordinates: 1,390,922–1,435,747) in *S. marcescens* N10A28 (GenBank: CP033623.1) following induction by mitomycin C. (L) Pre-colonization of *F. perrara* WT but not *F. perrara ΔclbB* induced the prophage-like region. Data are mean ± SD, *n* = 3 biological replicates. (M and N) Colibactin prodrug byproducts, *N*-lauryl-D-Asn (C12) and *N*-myristoyl-D-Asn (C14), measured by LC–MS in the ilea of gnotobiotic bees that were inoculated with *F. perrara* WT, *ΔclbB* or not inoculated. (O and P) Colibactin byproducts in the ileum of hive-collected bee foragers. The bees were sorted into “Scab” or “No Scab” groups based on pylorus phenotypes (Fig. S15). 20 ilea were pooled for each sample; data are mean ± SEM, *n* = 3 biological replicates in M–P. Statistical differences in L–P was assessed using a one-way ANOVA with a Tukey’s multiple comparisons test. Statistical significance: ****, *p* < 0.0001; ***, *p* < 0.001; **, *p* < 0.01; *, *p* < 0.05; ns, not significant.

In parallel, the plasmid pSL1-RCP, giving constitutive red fluorescence, was transferred to *F. perrara*. This setup allowed us to visualize the two bacteria and possible DNA damage in situ. *F. perrara* always colonized a short region from the pylorus to the anterior ileum, while *S. marcescens* was ubiquitous in the ileum when mono-inoculated (Fig. 2B–D). Pre-inoculation with *F. perrara* WT largely inhibited the proliferation of *S. marcescens* in the ileum (Fig. 2E–H), Occasionally, *S. marcescens* still formed aggregates with strong fluorescence intensity in the ileum (Fig. 2E), but in most cases, *S. marcescens* was either repelled from the anterior region (Fig. 2F) or was completely cleared from the ileum (Fig. 2G, H). In the presence of *F. perrara* WT, *S. marcescens* showed strong GFP signal indicating genotoxic activity. In contrast, *F. perrara ΔclbB* did not show a repellent effect towards *S. marcescens* (Fig. 2I). Measurements of mean fluorescence intensities of GFP and E2C in the ileum of different treatment groups showed that bees pre-colonized with *F. perrara* WT were higher in GFP and lower in E2C compared to *F. perrara ΔclbB* and MD (*p* < 0.001), which further corroborated that the protective effect is colibactin-dependent (Fig. 2J).

Silpe *et al.* reported that the genotoxicity of colibactin, as for other physical or chemical mutagens, triggers prophage induction (*29*). We next investigated whether *F. perrara* could induce prophages in the honeybee gut. *S. marcescens* strain N10A28 (Genbank: CP033623) has three prophage-like islands, of which two appear to have complete sets of phage genes (Supplementary Fig. S8). We first tested for prophage induction with MMC in vitro. By mapping short reads from Illumina sequencing to the genome, we observed that MMC treatment caused coverage for one prophage region to be 134.9-fold higher than coverage for other genomic regions, indicating that this prophage is inducible by genotoxicity (Fig. 2K). The other two prophage-like regions were not induced by MMC (Supplementary Fig. S9).

We then tested whether colibactin produced by *F. perrara* within the gut causes prophage induction in *S. marcescens*. Sequencing of gut samples indicated that coverage for the same *S. marcescens* prophage region was 6.4 times higher than coverage for other genomic regions in gut samples colonized by *F. perrara* WT, while coverage in this region was not elevated in guts colonized by *ΔclbB* (Fig. 2L and Supplementary Fig. S10). Thus, prophage induction resulting from *F. perrara-*produced colibactin is likely a mechanism underlying the observed inhibition of the pathogen.

### Colibactin byproducts in *F. perrara*-colonized bee guts

Next, we verified that *F. perrara* produces *clb*-dependent metabolites in vivo, as previously shown in vitro (*12*). Because direct quantitation of colibactin is still not possible owing to its instability, we used liquid chromatography–mass spectrometry (LC–MS) to detect two colibactin prodrug byproducts from *F. perrara*-colonized bee guts: *N*-lauryl-D-Asn (C_16_H_30_N_2_O_4_, C12 byproduct) and *N*-myristoyl-D-Asn (C_18_H_34_N_2_O_4_, C14 byproduct). They are the two most abundant *clb* metabolites identified in *F. perrara* pure culture (*12*). N-myristoyl-D-Asn is formed by NRPS ClbN in the first step of the biosynthetic pipeline in *E.coli* and cleaved from the biosynthetic precursor precolibactin by hydrolase ClbP in the last step, releasing the active colibactin genotoxin (*19, 20, 36–40*). We first assayed ilea (including pyloruses) of gnotobiotic bees inoculated with *F. perrara* WT, *ΔclbB* or not inoculated. After 10 days, levels of C12 and C14 byproducts in WT-colonized bees were 10.1- and 8.9-fold higher, respectively, than those in *ΔclbB*-colonized bees (Fig. 2M, N). In several other culture media, WT cell pellets had higher levels of C12 and C14 than *ΔclbB* (Supplementary Fig. S14), while in culture supernatants, the byproducts were below the detection limit.

To determine whether colibactin is being produced *by F. perrara* in outdoor hives, hive-collected bee foragers were sorted into two cohorts according to their pylorus “scab” phenotype (Supplementary Fig. S15), which indicates *F. perrara* colonization (*9*). Scab bees had 27.7- and 3.2-fold higher levels of C12 and C14 byproducts in ilea than no-scab bees (Fig. 2O, P), demonstrating the presence of *F. perrara*-produced colibactin.

Altogether, we detected high production of byproducts from *F. perrara* WT both in vivo and in vitro. Residual production of byproducts in *ΔclbB* can be attributed to assembly line derailment of the upstream ClbN step which remains intact in this mutant strain (*12*).

### Homologs of *clbS* for colibactin resistance are ubiquitous in honeybee specialists

In view of the negative effects of colibactin on *S. marcescens*, we wondered how colibactin production by *F. perrara* impacts members of the normal bee gut community. Colibactin producers, such as *F. perrara* and some *E. coli* strains, always encode a resistance gene, *clbS* (*41*), within the *clb* island (Fig. 1A). Some non-colibactin producers also have colibactin-resistance conferred by *clbS* (*29*), and we wondered whether non-colibactin-producing members of honeybee gut community possess this resistance mechanism. All up-to-date genomes from bacterial species isolated from honeybee guts or hives were collected and sorted into three categories: core symbionts (restricted to bee guts and present in all individuals), other bacteria restricted or not to bee guts, and pathogenic species known to affect bees. Surprisingly, Blastp results indicated that ClbS homologs (having 26–67% amino acid sequence identities to *E. coli* ClbS and 22–98% identities to *F. perrara* ClbS; Supplementary Data S1) were encoded in most (87-98%) genomes of species restricted to bee guts, including both Gram-negative and Gram-positive bacterial species (Table 1). Other than *F. perrara*, no other bee gut specialists encode the colibactin genomic island. By contrast, none of the pathogenic taxa, including *S. marcescens*, possessed *clbS*-like genes. Bacteria closely related to bee gut core taxa but living in other environments also rarely possess *clbS*, indicating likely acquisition through horizontal transfer by the bee gut bacteria (Supplementary Fig. S16 and Supplementary Data S1). Consistent with this scenario, the representative *clbS* genes appear in random positions within different genetic neighborhoods (Supplementary Fig. S17), and the *clbS*-based phylogeny shows considerable clustering reflecting bacterial taxonomy but also implying some transfer between distantly related bacteria, for example, between *Snodgrassella* and *Gilliamella* (Fig. 3A).

**Fig. 3.**
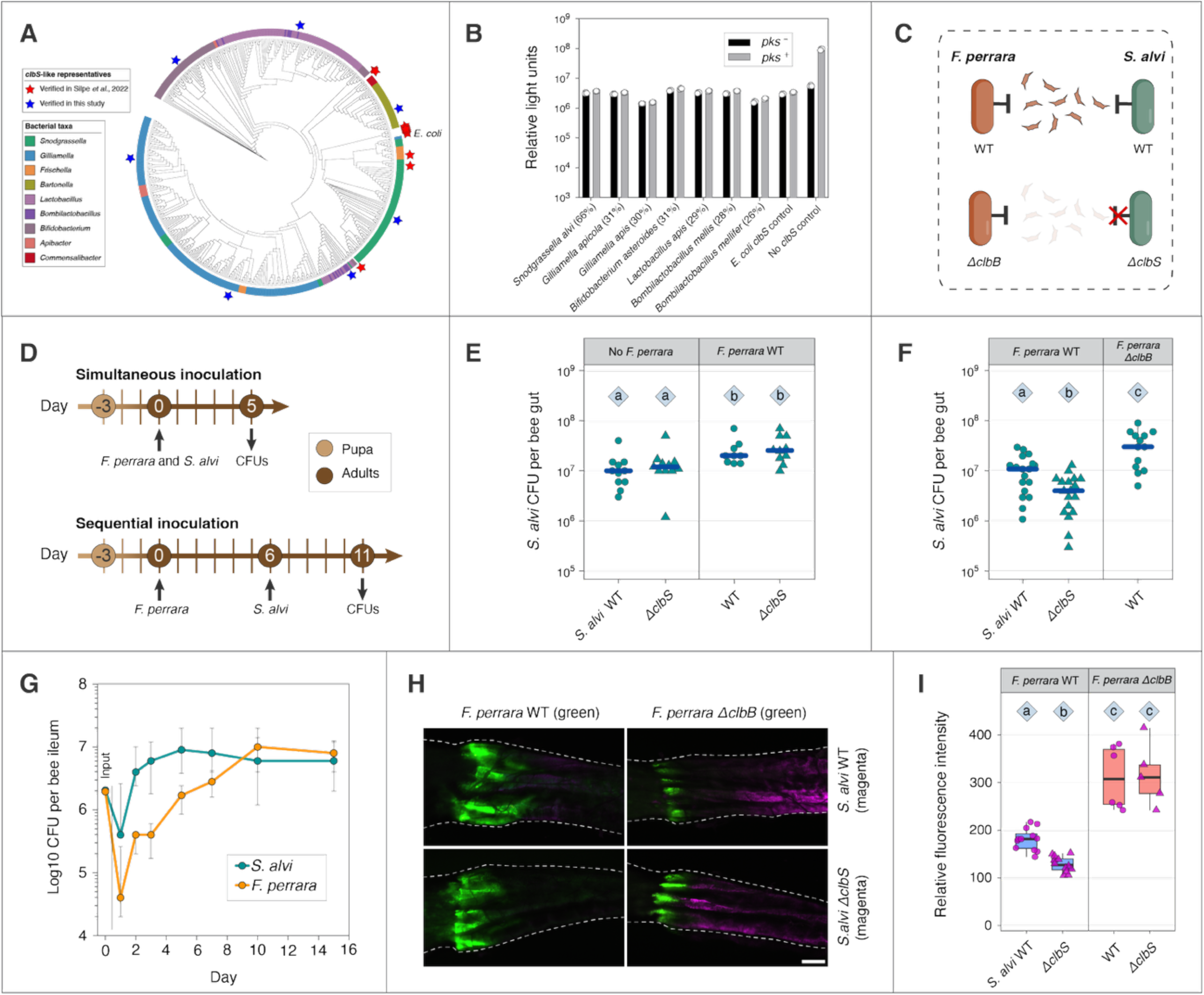
ClbS homologs from honeybee gut symbionts are protective against colibactin. (A) Phylogeny of all *clbS*-like genes from honeybee gut-related bacterial taxa. Representatives from human and bee guts (red pentagrams) were previously shown to protect *E. coli* from DNA damage and prophage induction caused by colibactin (*29*). The *E. coli* ClbS is labelled. Additional sequences representing different branches (blue pentagrams) were verified in this study. (B) A bioluminescent reporter assay showing that all honeybee gut *clbS*-like representatives are protective. Percent amino acid identity of each homolog to *E. coli* ClbS is given in parentheses. The DNA damage caused by *pks*^+^ *E. coli* increased light units when a functional *clbS* was absent. The presence of honeybee gut *clbS*-like genes reduced light units like the native *E. coli clbS*. Three biological replicates were measured for each treatment group. (C) Schematic representation of experimental strains. The T-shaped lines indicate the *clbS*-mediated resistance, and the red cross indicates that the gene deletion mutant lost this resistance. (D) Timelines of simultaneous and sequential inoculation of *F. perrara* and *S. alvi* to gnotobiotic bees. (E) Colonization of *S. alvi* strains in bee guts that were simultaneously inoculated with *F. perrara* or not inoculated (*χ^2^*= 15.8, *df* = 3, *p* = 0.0012, *n* = 41). (F) Colonization of *S. alvi* strains in bee guts that were pre-inoculated with *F. perrara* or not inoculated (*χ^2^* = 21.6, *df* = 2, *p* < 0.0001, *n* = 49). Blue lines show median values. (G) A time course measurement of the colonization of *F. perrara* and *S. alvi* in the ileum of gnotobiotic bees. The bees were inoculated with both species; 14-19 bees were measured per time point (total *n* = 120). Data are median and interquartile range. (H) Fluorescence microscopy showing the pylorus region of gnotobiotic bees colonized by *F. perrara* and *S. alvi* strains. *F. perrara* WT or *ΔclbB* were pre-colonized 6 days before inoculating *S. alvi* WT or *ΔclbS*, and guts were dissected and imaged 2 days after. Scale bar = 0.2 mm. (I) Fluorescence quantification of *S. alvi* strains in bee guts pre-colonized with *F. perrara* WT or *ΔclbB* (*χ^2^* = 29.6, *df* = 3, *p* < 0.0001, *n* = 35). For (E), (F), and (I), statistical significance was assessed using Kruskal–Wallis test (reported in parentheses for each experiment), followed by post-hoc pairwise comparisons of treatment groups using Dunn’s test. Groups with different letters are significantly different (*α* = 0.05).

**Table 1.** Presence of *clbS*-like genes among honeybee gut-related bacterial taxa.*

|  | Number of<br>genomes with<br>*clbS* | Total number<br>of genomes | Percent of genomes<br>with *clbS* |
| --- | --- | --- | --- |
| **Core gut symbionts** |  |  |  |
| *Snodgrassella alvi* | 66 | 68 | 97 |
| *Gilliamella* spp. | 161 | 177 | 91 |
| *Bifidobacterium* spp. | 62 | 63 | 98 |
| *Lactobacillus* spp. | 105 | 115 | 91 |
| *Bombilactobacillus* spp. | 13 | 15 | 87 |
| **Non-core, restricted to bee gut** |  |  |  |
| *Frischella* spp. | 13 | 16 | 81 |
| *Bartonella* spp. | 31 | 31 | 100 |
| **Not bee-gut restricted** |  |  |  |
| *Apibacter* spp. | 10 | 29 | 34 |
| *Commensalibacter* spp. | 6 | 43 | 14 |
| *Apilactobacillus* spp. | 0 | 147 | 0 |
| *Bombella* spp. | 0 | 20 | 0 |
| *Parasaccharibacter* spp. | 0 | 14 | 0 |
| **Pathogens** |  |  |  |
| *Serratia marcescens* | 0 | 5 | 0 |
| *Hafnia alvei* | 0 | 2 | 0 |
| *Melissococcus plutonius* | 0 | 27 | 0 |
| *Paenibacillus larvae* | 0 | 40 | 0 |

\*Genomes and sequences are detailed in Supplementary Data S1.

Silpe *et al.* showed that *clbS*-like genes from non-colibactin-producing bacterial species, including *S. alvi*, provide protection from colibactin-induced DNA damage and prophage induction (*29*). Using a similar assay, we tested *clbS* homologs from representatives of all core members of the bee gut community. Briefly, *clbS* homologs were cloned into an *E coli* strain that reports DNA damage as bioluminescence, and protection was tested by co-cultivating this strain with either a *pks*+ or *pks*-*E. coli* strain. All tested *clbS* homologs provided protection against colibactin-induced DNA damage (Fig. 3B). Thus, core community members may be subject to chronic colibactin-based selective pressure, leading to acquisition and maintenance of the protective *clbS*.

### ClbS homolog protects *S. alvi* from colibactin genotoxicity

We tested the protective effect of *S. alvi clbS* in vivo using a *clbS* deletion mutant (Fig. 3C). First, we simultaneously inoculated either *S. alvi* WT or *ΔclbS* with and without *F. perrara* WT and evaluated CFUs after 5 days (Fig. 3D). Surprisingly, whether *F. perrara* WT was co-inoculated or not, *S. alvi* WT and *ΔclbS* had similar levels of colonization (*p* = 0.86, 0.56, respectively) (Fig. 3E). Interestingly, both *S. alvi* strains were more abundant in guts pre-colonized with *F. perrara* WT (*p* < 0.05).

We then sequentially inoculated bees with the two species (Fig. 3D). When *F. perrara* WT was established first, *S. alvi* WT colonized 2.5-fold higher than *S. alvi ΔclbS* (*p* = 0.019) (Fig. 3F). When *F. perrara ΔclbB* was established first, *S. alvi* WT colonized 2.4-fold higher than when *F. perrara* WT was established first (*p* = 0.020).

To explore the basis of the delay in effect of *F. perrara* colonization on *S. alvi*, we performed a time-course measurement of colonization by the two species. With the same amount of inocula (∼2×10^6^ cells), *S. alvi* reached a maximum of ∼10^7^ cells in the ileum after 2 days, while *F. perrara* slowly increased to ∼10^7^ cells after 10 days (Fig. 3G). The delay is likely due to the slower establishment of *F. perrara*.

Using strains carrying constitutive GFP/E2C expression plasmids, we observed the influence of pre-colonized *F. perrara* on *S. alvi* colonization in the anterior hindgut. When *F. perrara* WT was pre-colonized, *S. alvi ΔclbS* showed weaker colonization compared to *S. alvi* WT after 2 days (*p* = 0.012) (Fig. 3H, I). In contrast, when *F. perrara ΔclbB* pre-colonized, *S. alvi* WT and *S. alvi ΔclbS* colonized to similar levels (*p* = 1.0) which were higher than when *F. perrara* WT pre-colonized (*p* = 0.033, < 0.0001, respectively). These results reveal that *F. perrara* exerts both enhancing and colibactin-dependent inhibitory effects on *S. alvi* and that *clbS* confers protection against colibactin in *S. alvi*.

We also evaluated the influence of *F. perrara* colonization on other gut bacterial members with a defined community of representatives of core bee gut lineages (see Supplementary Text). Except for *Gilliamella,* other taxa are not significantly reduced in the hindgut even when *F. perrara* is pre-established. Community composition and size exhibit no effects from colibactin pressure. Based on our results, a likely reason is that all the included strains possess *clbS*.

## Discussion

### Evolutionary consequences of the intrusion of a colibactin producer in a natural community

Many animals host a characteristic gut microbiota, the core members of which may have undergone long-term coevolution with their hosts and with one another (*42–44*). Microbial warfare appears to be pivotal in these communities, driven by competition for spatial niches and nutrients. Broad-spectrum toxins such as colibactin can have especially potent effects on communities.

Some studies in mice (*28, 32*) found that colibactin production influences gut community structure. However, the mechanisms of action of colibactin within natural communities and its impacts on longer term evolution of community members are largely unknown. Using the simple natural community of the honeybee, we found that this broad-spectrum genotoxin produced by a gut symbiont excludes pathogens while having limited effects on the normal community, thereby benefiting the host (Fig. 4). Our in vivo results provide direct evidence that colibactin causes DNA damage and prophage induction in situ and thus support its role as a weapon produced for interbacterial competition in nature. The ubiquity of the resistance genes among specialists in this system brings new evolutionary insights into how bacterial weapons can shape the evolution and ecology of natural communities.

**Fig. 4.**
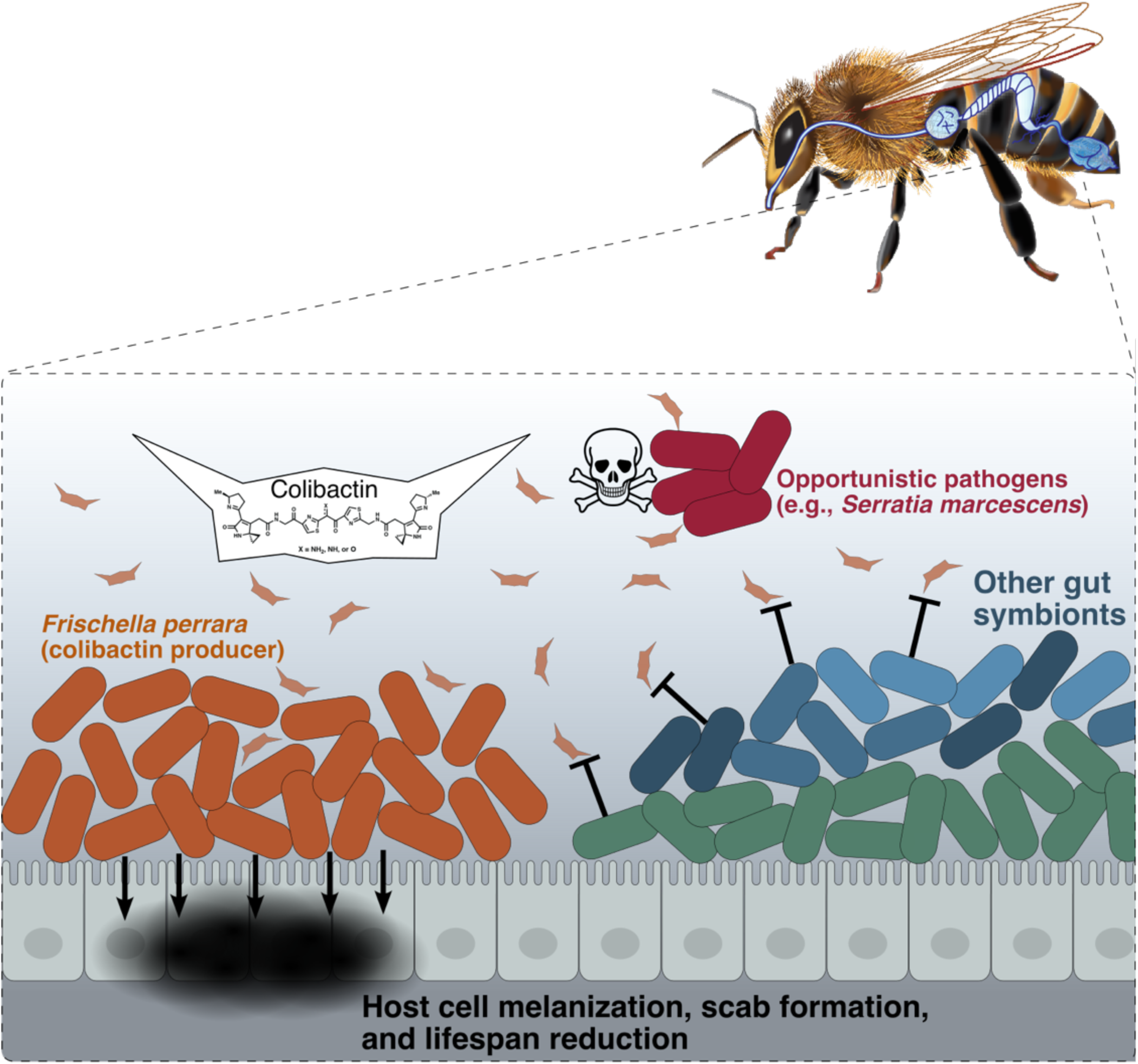
Schematic representation of influences of *F. perrara* colonization and colibactin production on other bacteria and the honeybee host. *F. perrara* guards the entrance of the hindgut where the microbial community are colonizing. Colibactin production by *F. perrara* generates selective pressure on community members, leading to ubiquitous resistance of the core members. The T-shaped lines indicate *clbS*-mediated resistance. Pathogens that lack the resistant mechanism are excluded, benefitting the host. In the absence of pathogens, *F. perrara* colonization potentially increases bee mortality due to genotoxicity of colibactin and overactivation of host immune responses.

*F. perrara* and other species in the same genus exclusively colonize guts of honeybees (*Apis* species) (*6*). Unlike other major bee gut taxa, which are associated with all major lineages of corbiculate bees and which were established in a common ancestor, *Frischella* has a more recent evolutionary association with bees and is restricted to the honeybee branch (*44*). *F. perrara* produces colibactin, which causes DNA damage and can induce prophages ((*12*) and this study).

Our results provide a spatial context for effects of colibactin on a gut community. We observed that *F. perrara*-produced colibactin resulted in repulsion or complete exclusion of *S. marcescens* from the gut ileum. Abdomens of *F. perrara*-colonized bees had lower *S. marcescens* titers overall, suggesting that this suppression extends throughout the hindgut. In addition, the ubiquity of functional *clbS* homologs in core members of the bee gut community supports the hypothesis that colibactin exerts ongoing selective pressure throughout the honeybee hindgut. Even Gram-positive species (*Lactobacillaceae* and *Bifidobacterium*), that dominate in the rectum, usually possess *clbS*.

*clbS* encodes a DNA-binding putative cyclopropane hydrolase that imparts colibactin resistance (*41, 45, 46*). Protection by ClbS appears to be primarily intracellular, as supernatants from ClbS-expressing bacterial cells do not protect against colibactin (*29*). However, extracellular supplementation with ClbS attenuates colibactin genotoxicity to cultured human cells (*18, 47*), suggesting the possibility that ClbS released from dead cells may confer protection to surrounding cells. However, the ubiquity of *clbS* homologs in bee gut taxa indicates that native expression of ClbS homologs provides an evolutionary advantage.

### Gut microbial interactions revealed by gnotobiotic bee assays

Our study illustrates the complexity of microbial interactions even in a relatively simple community. In the gut, *F. perrara* establishes much more slowly than *S. alvi* and other core symbionts. It inhibits *S. alvi* when preinoculated but not when coinoculated, which suggests that the impact of colibactin may depend on cell density or sufficient time for pathway induction.

*F. perrara* boosts colonization by *S. alvi*, especially when it does not produce colibactin. *F. perrara* is an obligate anaerobe with a sugar-fermenting metabolism, whereas *S. alvi* is an aerobe that does not directly utilize sugars but oxidizes short-chain fatty acids. *S. alvi* likely benefits from the fermentation products produced by *F. perrara*, as has been proposed for *Gilliamella* spp. which also ferment sugars (*7*).

*Gilliamella apicola* and *Gilliamella apis* are fermentative bacteria adjacent to *F. perrara* in the ileum. Our analyses show that their levels are negatively correlated to those of *F. perrara*, both in defined community experiments in gnotobiotic bees and in hive bees, suggesting competition between the two genera in the gut. The inhibitory effect of *F. perrara* on *Gilliamella* spp. is colibactin-independent, since the effect occurs even for *F. perrara ΔclbB*. Other mechanisms, e.g., type VI secretion systems (*48*), could contribute to the negative correlation.

The advantages of colibactin production to *F. perrara* may reflect a metabolic advantage in the host niche, a competitive advantage within the community, and/or an effect on host cells that favors *F. perrara* expansion. *F. perrara ΔclbB* colonization is weaker than that of WT in mono-colonization or in presence of other community members ((*17*) and this study). Likewise, *pks^+^ E. coli* has advantages over *pks^-^*counterparts in gut colonization (*33, 49*), possibly because genes of this locus are also implicated in production of other metabolites such as γ-lactam derivatives (*50*), siderophores (*51*) and microcin (*52*), which may contribute to colonization capacity. It is not clear whether *clb* genes in *F. perrara* are similarly linked to such compounds.

### Effect of oxygen on colibactin persistence

In vitro studies of colibactin-producing *E. coli* (*pks^+^*) suggested that cell-cell contact is required for genotoxicity as *pks^+^*supernatant or physical separation of *pks^+^* cells from eukaryotic or bacterial cells did not result in toxicity (*13, 29*). However, a more recent study indicates that the effect is contact-independent as DNA damage occurs when producer and reporter colonies are spatially separated (*30*). These different results most likely reflect the instability of colibactin. Biosynthetic and structural studies suggest that mature colibactin contains a reactive α-aminoketone structural motif that is susceptible to aerobic oxidation (the α-ketoimine is found in an NMR structure of the colibactin interstrand crosslink) and hydrolysis, ultimately leading to fragmentation of the compound (*20, 21, 40, 53, 54*). In *pks^+^ E. coli*, the production of colibactin is maximal under anoxic conditions and decreases with increased oxygen concentration, and the DNA cross-linking activity drops sharply with increasing oxygen levels and becomes undetectable at atmospheric oxygen, which suggest that the regulation of the *pks* genes and potential chemical decomposition of colibactin are relevant to oxygen status (*55*). Anoxia is optimal for the activity and production of colibactin, which may explain the prevalence of *pks^+^ Enterobacteriaceae* strains in the human gut environment where there is a steep gradient between the hypoxic epithelium and anoxic lumen (*55*). In the tiny honeybee gut, the epithelial bacterial biofilm actively consumes oxygen, forming a steep gradient in the ileum, the central region of which is completely anoxic (*56*). Presumably, the low oxygen environment enables a longer half-life and considerable activity of the toxin. Colibactin is probably also present in the rectum which possesses more anoxic volume.

### Consequences for host health

*F. perrara* may benefit bee hosts by excluding pathogens. Chen *et al.* (*28*) reported that a commensal *E. coli* inhibits human pathogens including *Vibrio cholerae* and *Bacteroides fragilis* in a colibactin-dependent manner both in vitro and in mouse intestines. Interestingly, their analysis on murine and human gut microbiome datasets revealed that *pks*+ *E. coli* reduces *B. fragilis* but no other commensal *Bacteroides* spp. Similarly, the different outcomes of different strains might be explained by the presence of ClbS homologs. Though Silpe *et al.* (*29*) found ClbS homologs in non-colibactin-producing members, their presence in major taxa of the human gut microbiome has not been systematically evaluated. *E. coli* Nissle 1917 requires colibactin production for its probiotic activity (*57*). A role of colibactin in excluding alien strains including pathogens may be a general phenomenon in different gut systems, including both bees and humans.

On its own, *F. perrara* is detrimental to bee health. Though the scab-like phenotype caused by *F. perrara* is reminiscent of the link between colibactin-producing *E. coli* and colorectal cancer (*25, 26, 58*), scab-formation is colibactin-independent (*17*). Scabs likely result from innate immune stimulation leading to the melanization cascade (*11*), potentially mediated by a T6SS (*17*). Overactivated immune responses might contribute to the observed increase in mortality (Supplementary Fig. S3). Genotoxicity may also contribute to observed negative effects of colibactin production on host survivorship.

### Conclusions

The honeybee gut system provides a simple animal model for identifying mechanisms underlying coevolved interactions. Using this system, we elucidated the impacts of a relatively recently evolved symbiont and its genotoxin on a gut community and its dual effects on host health. We found that community members have acquired resistance genes that lessen the negative impact of the toxin, supporting the hypothesis that colibactin is produced as a weapon for interbacterial competition in gut environments. Because non-members, including pathogens, lack these protective genes, this genotoxin serves to preserve community composition, with beneficial outcomes for hosts. Our findings shed light on the intricate interactions of microbial members with one another and with the host.

## Materials and Methods

### Bacterial strains, plasmids, and primers

The bacterial strains used in this study are listed in Table S1; plasmids are in Table S2; primers are in Table S3.

### Bacterial culture

*Frischella perrara*, *Snodgrassella alvi*, *Serratia marcescens*, and *Bifidobacterium* and *Gilliamella* strains were cultured on Columbia agar (BD, Franklin Lakes, NJ, USA) supplemented with 5% sheep blood (HemoStat Laboratories, Dixon, CA, USA), with or without specified antibiotics. *Lactobacillus* and *Bombilactobacillus* strains were cultured on Man, Rogosa, and Sharpe (MRS; HiMedia Laboratories, Maharashtra, India) agar. *F. perrara* was grown at 35 °C in an anaerobic chamber; other bee gut isolates were in 35 °C incubator with 5% CO_2_.

Per experiment, bacterial colonies were inoculated into Columbia broth (CB; BD) or MRS and grown statically in 35 °C incubator with 5% CO_2_, except *F. perrara* was grown in rubber-stopped anaerobic Hungate tubes with a 100% N_2_ headspace.

*E. coli* was cultured on Lysogeny broth (LB) agar at 37 °C or in LB with orbital shaking at 200 rpm at 37 °C. 0.3 mM diaminopimelic acid (DAP) was supplemented for *E. coli* MFD*pir*.

The following antibiotic concentrations were added as needed: ampicillin (100 µg/mL *E. coli*, 50 µg/mL *S. alvi*), kanamycin (50 µg/mL *E. coli*, 25 µg/mL *S. alvi*, 25 µg/mL *F. perrara*, 200 µg/mL *S. marcescens*).

### Gene deletion in *S. alvi*

*S. alvi clbS* gene was knocked out by direct transfer of a recombineering cassette through electroporation, as previously described (*59*). Q5 High-Fidelity DNA Polymerase (NEB, Ipswich, MA, USA) were used for PCR reactions. Annealing temperatures were as recommended by NEB Tm calculator (https://tmcalculator.neb.com/). PCR products were collected with AxyPrep Mag PCR Clean-up kit (Axygen Biosciences, Union City, CA, USA).

The gene deletion cassette was generated via a three-step assembly. First, homology arms (1200-1400 bp) flanking the gene coding region were amplified using genomic DNA as template, and the ampicillin resistant cassette (*ampR*) was amplified using pBTK501 (*60*) as template. Second, about 200 ng each of the collected 5’ and 3’ arm fragments and *ampR* were pooled in a 25 µL fusion reaction (with other PCR components but no primers) for 30 cycles. Last, 1 µL of the fusion output was used as template for a following PCR, with the forward primer of 5’ arm and the reverse primer of 3’ arm, to generate enough amount of the assembled cassette for transfer.

Ampicillin-resistant transformants were checked by PCR using a pair of screen primers and further verified by submitting to Plasmidsaurus (https://plasmidsaurus.com/) for whole genome sequencing.

### Bioluminescent reporter assay for *clbS*-like genes

Protection by honeybee gut *clbS*-like genes against colibactin was tested in vitro via a bioluminescent reporter method as previously reported (*29*). Open reading frames (ORFs) of selected genes were codon-optimized for *E. coli* expression using Codon Optimization Tool from Integrated DNA Technologies (IDT; https://www.idtdna.com/pages/tools/codon-optimization-tool), and the optimized sequences (Table S4) were synthesized and cloned into pTrc backbone by Twist Bioscience (Twist Bioscience, South San Francisco, CA, USA). The plasmids were transformed into *E. coli* BW25113 harboring the P_R_-*lux* reporter. The full-length WT *clbS* from *E. coli* CFT073 and an empty vector were used as positive and negative controls, respectively. Strains were grown overnight in LB (RPI, Mt. Prospect, IL, USA) at 37 °C, then back-diluted 1:100 and co-cultured in M9+CAS medium (Quality Biological, Gaithersburg, MD, USA) at a 1:1 ratio with either *E. coli* BW25113 harboring BAC-*pks* or *E. coli* BW25113 harboring an empty vector. Luminescence was measured after 20 h of incubation at 37 °C in a BioTek Neo2 multi-mode plate-reader (Bio Tek, Winooski, Vermont, USA) and readings were normalized by endpoint OD_600_.

### Plasmid construction and transformation

Fluorescent reporter cassettes were designed to report DNA damage in *S. alvi* wkB2 and *S. marcescens* N10A28, respectively. The cassettes comprise a promoter region (100-200 bp upstream the *recA* gene homolog of wkB2 or N10A28), eGFP gene, and a terminator (Table S4). DNA sequences were synthesized by Twist Bioscience. The plasmid backbone of pSL1-E2C and the fluorescent cassettes were amplified, digested with BsaI or BsmBI, and ligated with T4 DNA ligase (Promega, Madison, WI, USA). The ligation products were transferred to *E. coli* DH5α (NEB). Constructed plasmids were sequenced in Plasmidsaurus.

Plasmids were conjugated to recipient strains via MFP*pir* donor as previous described (*60*).

### Fluorescence flow cytometry

*S. marcescens* carrying different fluorescent plasmids and WT were inoculated to 5 mL Columbia broth, with or without kanamycin. 5 µL overnight culture was inoculated to 1 mL fresh medium and grew with mild shake (∼50 rpm) for 5 hours; 0.05 µg/mL or 0.5 µg/mL mitomycin C (MMC) was added for induction. Cells were collected by centrifugation, washed twice and resuspended in PBS to ∼0.1 at OD_600_ that measured by a BioSpectrometer (Eppendorf, Enfield, CT, USA). Flow cytometry was conducted on a Fortessa Flow Cytometer (BD). About 10000 events were measured for each sample. Data was analyzed and visualized using FlowJo (v10; https://www.flowjo.com/).

### Preparation of microbiota-deprived honeybees and defined inoculation

We harvested late-stage pupae of *Apis mellifera* workers from brood frames within hives located at the University of Texas at Austin by carefully removing the surface wax with clean forceps. Pulled pupae were transferred to sterile plastic containers with a layer of absorbent bench underpads (VWR, Radnor, PA, USA) taped to the bottom and kept at 35 °C and approximately 60% relative humidity. Sterile 1:1 sucrose:water (*w/v*) solution (hereafter referred to as sucrose water) and irradiated pollen were supplied in the containers. After 3 days, the newly emerged adult bees, which lack their native microbiota (*34*), received bacterial inoculation by oral feeding.

Bacterial cells were collected from liquid cultures in late exponential phase (typically 2-3 days for the symbionts and overnight for *S. marcescens*) at 6,000 g x 5min, 4 °C and resuspended in PBS (pH 7.4). Unless otherwise mentioned, cell suspensions were always diluted to OD_600_ of 1 for bee inoculation. Antibiotics were not added to bacterial cultures or sucrose water unless otherwise indicated (for plasmid-containing strains).

The clean bees were starved for 2–3 hours, immobilized with CO_2_, and placed in 0.5 mL capped microcentrifuge tubes (tip removed) for feeding. Bacterial cell suspensions were aliquoted to 100 µL and mixed with 25 µL sucrose water (*v/v*, 80:20) immediately before feeding 5 µL per strain to each bee (to avoid that bacterial cells are killed by high sucrose osmolarity). The inoculated bees were put in clean cup cages (10-20 bees per cage) that were supplied with sucrose water and irradiated pollen and kept in a 35 °C incubator. For sequential inoculation or pathogen challenge, caged bees were immobilized with CO_2_ after 5 or 6 days and fed again in the same way.

### Survival and CFU assays of gnotobiotic honeybees

Newly emerged bees were inoculated with *F. perrara* WT/*ΔclbB* as above described. Uninoculated microbiota-deprived (MD) controls received PBS. For pathogen challenge, each bee was fed with 5 µL of *S. marcescens* (OD_600_ of 1) with sucrose water (*v/v*, 80:20; control groups received 5 µL PBS) after 5 days. Dead bees were counted and removed from cup cages at a similar time each day. Bees that died at day 1 were excluded (censoring) as they most likely died from the handling.

In the CFU assay, a sublethal density (OD_600_ of 0.5) of *S. marcescens* was fed to bees 5 days after *F. perrara* inoculation, and bees were dissected after 1 day and 3 days for CFUs. *S. alvi* WT/*ΔclbS* was fed to the bees together with *F. perrara* at day 0 (simultaneous inoculation) or after 6 days (sequential inoculation), and bees were dissected after 5 days for CFUs. Bee ilea (including pyloruses), hindguts, or abdomens were homogenized in 100 µL PBS for 15 s using a pestle BioVortexer (Biospec Products, Bartlesville, OK, USA). Serial dilutions (10 x) in PBS were spotted onto agar plates and CFUs were counted after 3 days of cultivation (1 day for *S. marcescens*).

### ROS assay

ROS activity determination was modified from Guo *et al.*, 2023 (*61*). Gnotobiotic bees inoculated with different bacteria were kept in cup cages supplemented with sucrose water but no pollen (as the autofluorescence from pollen interferes with subsequent determination). After 6 days, bees were dissected in a catalase inhibitor solution (20 mM 3-amino-1,2,4-triazole (TCI, Tokyo, Japan) in PBS). Malpighian tubules and other tissues were removed from the gut. Hindguts were homogenized in 100 µL inhibitor solution and centrifuged (12,000 *g* x 10 min, 4 °C). Hydroperoxides were quantified by Pierce™ Quantitative Peroxide Assay Kits (Thermo Fisher Scientific, Waltham, MA, USA). 20 µL supernatant was used for each reaction (x 3 technical replicates for each sample). Absorbance was measured at 595 nm by a Tecan Spark 10M plate reader (Tecan Group, Morrisville, NC, USA). A calibration was made with hydrogen peroxide for the quantification according to the manual.

### Inoculation of a defined bacterial community and high throughput sequencing

Newly emerged bees were inoculated with *F. perrara* WT, *ΔclbB*, or not inoculated (MD). The bees were kept individually in 50 mm petri dishes supplemented with sucrose water and irradiated pollen (*62*). A defined community comprising 9 strains (Table S1) was inoculated after 5 days. All strains were resuspended together at OD_600_ of 1 for each strain. This bacterial suspension was mixed with sucrose water (*v/v*, 80:20), and 5 µL was fed to each bee. Bees were dissected after another 5 days, and DNA was extracted from hindgut of each individual bee using PureLink™ Microbiome DNA Purification Kit (Thermo Fisher Scientific).

We produced metabarcoded amplicons of the V4 region of 16S rRNA genes for high throughput sequencing, via a two-step amplification strategy similar to that in Powell *et al.*, 2021 (*35*). The first PCR was in 25 µL reactions that contain 1 µL diluted template DNA (∼10 ng), 1 µL of 10 µM Hyb515F_rRNA and Hyb806R_rRNA (Table S3), 12.5 µL 2x Accustart II PCR Supermix (Quantabio, Beverly, MA, USA), and 9.5 µL molecular grade water. Cycling conditions were 94 °C for 3 min; 30 cycles of 94 °C for 20 s, 50 °C for 15 s, 72 °C for 30 s; followed by 72 °C for 10 min. We examined amplicons on a 2% agarose then purified with 0.8x HighPrep™ PCR magnetic beads (MAGBIO, Gaithersburg, MD, USA). The cleaned products were recovered in a final volume of 50 µL. The second PCR attached Illumina 8 bp Nextera style dual-indexed barcodes to products of the first PCR using unique combinations of N7XX and S5XX barcodes. It was performed in 25 µL reactions that contain 5 µL template DNA (cleaned from PCR 1), 2 µL of 5 µM indexed Hyb_Fnn_i5 and Hyb_Rnn_i7 (Table S3), 12.5 µL Accustart II Supermix, and 3.5 µL molecular grade water. Cycling conditions were 94 °C for 3 min; 10 cycles of 94 °C for 20 s, 55 °C for 15 s, 72 °C for 60 s; followed by 72 °C for 10 min. These reactions were cleaned with magnetic beads, resuspended in 25 µL molecular grade water and quantified using Qubit™ dsDNA Broad Range Quantitation Kit and a Qubit 4 Fluorometer (Thermo Fisher Scientific). We pooled equimolar amounts of each amplification, combined with 7% phiX DNA and sequenced on an Illumina iSeq 100 system (model number: FS10000184, Illumina Inc., San Diego, CA, USA) at 2 x 150 reads.

The amplicon sequencing data were processed, analyzed and visualized with QIIME 2 (v2023.9) (*63*), following the “Moving Picture” tutorial (https://docs.qiime2.org/2023.9/tutorials/moving-pictures/). Primer sequences were removed using the cutadapt plugin (*64*). Truncated reads were demultiplexed, quality filtered, followed by denoising with the Deblur plugin (*65*). Taxonomy was as198signed to amplicon sequence variants (ASVs) based on the Silva database (v138_Ref_NR_99) (*66*) using feature-classifier plugin (*67*). Unclassified and contaminant (mitochondria and chloroplast) ASVs were filtered out. An ASV table was generated for community profiling.

Total copies of 16S rRNA genes in each DNA sample was estimated via quantitative PCR as reported in previous studies (*68, 69*). The qPCR was performed in 10 µL reactions (triplicates for each sample) that contained 1 µL template DNA (10 x diluted), 0.6 µL of 5 µM 27F/355R (Table S3) universal primers for bacterial 16S genes, 5 µL 2x Bio-Rad SYBR Green master mix (Bio-Rad Laboratories, Hercules, CA, USA), and 3.4 µL molecular grade water, using 384 well format on an Applied Biosystems Quantstudio 5 Real-Time PCR system (Thermo Fisher Scientific). Cycling conditions were 95 °C for 10 min; 5 cycles of 95 °C for 15 s, 65-60 °C (−1 °C/cycle) for 15 s, 68 °C for 20 s; 35 cycles of 95 °C for 15 s, 60 °C for 15 s, 68 °C for 20 s. A serial dilution of plasmid was used as standard.

### Fluorescence microscopy

Bacterial strains carrying fluorescent plasmids were grown with kanamycin and inoculated into gnotobiotic bees as aforementioned. *F. perrara* was fed at day 0 and *S. alvi* was fed after 6 days. 25 µg/mL kanamycin was added to the bacterial cell suspension during feeding and to the sucrose water in cup cages pollen was not supplemented). Fluorescence in the pylorus and ileum was inspected after another 5 days using a Nikon AXR Confocal Microscope. (Our fluorescence method does not allow us to directly examine the rectum due to its low translucency and autofluorescence from gut contents.) *S. marcescens* was fed at OD_600_ of 0.5 after 6 days of *F. perrara* pre-inoculation. Kanamycin concentration in the sucrose water was increased to 100 µg/mL after feeding *S. marcescens*. After another 2 days, fluorescence was inspected using a Nikon Eclipse TE2000-U Inverted Fluorescence Microscope (Nikon Instruments, Melville, NY, USA). Fluorescence intensities were quantified using NIS-Elements AR (v5.42.03).

### Prophage induction in vitro and in vivo

*S. marcescens* was grown overnight and *S. alvi* was grown for 2 days in CB. Cells were harvested by centrifugation and resuspended in fresh CB to ∼1 at OD_600_. 0.5 µg/mL MMC was added and induced for 5 hours. Cell pellets were collected by centrifugation.

Newly emerged bees were inoculated with *F. perrara* WT, *ΔclbB*, or not inoculated (MD). After 6 days, each bee was fed with 5 µl *S. marcescens* (OD_600_ of 0.5) or *S. alvi ΔclbS* (OD_600_ of 1) mixed with sucrose water (*v/v*, 80:20). *S. marcescens*-fed bees were dissected after ∼36 hours, and *S. alvi*-inoculated bees were after 4 days. Four hindguts were pooled into each sample.

DNA was extracted using PureLink™ Microbiome DNA Purification Kit (Thermo Fisher Scientific). DNA samples were sent to SeqCenter (https://www.seqcenter.com/) for Illumina sequencing. 200 Mbp (1.33M reads) and 1Gbp (6.67M reads) packages were used for pure cultures and gut samples, respectively.

The short-read sequencing data were aligned to reference genomes (GenBank: *S. marcescens* N10A28, CP033623.1; *S. alvi* wkB2, CP007446.1) and coverages to genomic regions were estimated using *breseq* (v0.38.3) (*70*). Statistical significance was analyzed by a one-way ANOVA with a Tukey’s multiple comparisons test (Graphpad Prism).

### Quantitation of colibactin prodrugs

For bee gut samples, 20 ilea were pooled into an Eppendorf tube and flash frozen. The samples were thawed on ice, and 400 µL of extraction solution (MeOH:MeCN:H_2_O, 2:2:1 ratio, containing 100 nM of D-27 myristoyl-prodrug internal standard) was added to each tube. Each sample was homogenized using a pestle homogenizer (VWR) for 15 seconds. Homogenized contents were transferred to bead-bashing tubes (Bio-Rad), and bead-beat twice for 45s at room temperature, with cooling on ice after each agitation (approx. 5 minutes). Samples were centrifuged (5,000 *g* x 5 min, 4 °C), and supernatants transferred to a separate microcentrifuge tube and spun again (21,300 *g* x 20 min, 4 °C). The supernatant was carefully aspirated then filtered through a 0.2 µm nylon filter (Cytiva, Marlborough, MA, USA) into an autosampler vial. Supernatants were analyzed by UPLC–MS/MS on a Waters Xevo TQ-S UPLC-triple quadrupole with a Acquity UPLC H-Class System using a Cortecs UPLC C8 column (1.6µm, 2.1 mm x 75 mm) (Waters Corporation, Milford, MA, USA). The conditions were as follows: 0.5 mL/min flow rate, column temperature: 40 °C, 1 µL injection, 10% solvent B in solvent A for 0.5 min, a linear gradient increasing to 95% solvent B in solvent A over 0.5 min, holding 95% solvent B in solvent A for 1.0 min, followed by a linear gradient back to 10% solvent B in solvent A over 0.6 min, and re-equilibration at 10% solvent B in solvent A for 0.9 min (Solvent A, water + 0.1% formic acid; Solvent B, acetonitrile + 0.1% formic acid). The mass spectrometer was run in negative mode MRM, with the following conditions: capillary voltage, 2.1 kV, cone voltage, 4 V; source offset voltage, 50 V; desolvation temperature, 200 °C; desolvation gas flow, 800 L/h; cone gas flow, 150 L/h; Nebulizer, 7.0 ba. Ions of interest were quantified by monitoring the transitions *m/z* 313.2 → *m/z* 198.3 (cone voltage 50 V, collision energy 24 V, retention time 1.82 min) for *N-*lauryl-D*-*Asn, *m/z* 341.3 → *m/z* 226.3 (cone voltage 50 V, collision energy 24 V, retention time 1.90 min) for *N-*myristoyl-D*-*Asn, and *m/z* 368.5 → *m/z* 253.3 (cone voltage 58 V, collision energy 28 V, retention time 1.91 min) for the deuterated *N-*myristoyl-D*-*Asn internal standard. Each sample was collected with two replicate injections, with three independent triplicates. The ion counts of *N-*lauryl-D*-*Asn in each sample injection were normalized to the ion counts of the deuterated-*N-*myristoyl-D*-*Asn internal standard. A calibration curve of an authentic standard of *N-*myristoyl-D*-*Asn (0 – 500 nM) was used to find the concentration of *N-*myristoyl-D*-*Asn in samples, and the concentration of *N-*myristoyl-D*-*Asn was normalized to the concentration of the deuterated-*N-*myristoyl-D*-*Asn internal standard (100 nM).

For pure cultures, *F. perrara* WT and *ΔclbB* were inoculated to 50 mL Columbia broth or LB. After 2 days of incubation at 35°C in an anaerobic chamber, cell pellets were collected from CB cultures at 6,000 *g* x 5 min, washed twice with autoclaved water, and transferred into either 20 mL CB or 20 mL M9 medium (amended with 0.1% yeast extract and 0.1% Casamino acids). The transferred CB and M9 cultures were incubated for another 16 hours in the anaerobic chamber. The LB cultures were directly grown for three days (without a transfer). Cell pellets and supernatants were collected at 6,000 *g* x 5 min and flash frozen. Cell pellet samples were thawed on ice, added with 400 µL of aforementioned extraction solution containing an internal standard, sonicated in a water bath for 5 minutes (VWR), and further processed in the same way as bee gut samples. For the culture supernatants, 300 μl was lyophilized overnight, extracted with 600 μl extraction solution and further processed in the same way.

Samples were analyzed by a two-tailed unpaired t-test or analyzed by a one-way ANOVA with a Tukey’s multiple comparisons test for statistical significance (Graphpad Prism).

### Bioinformatics

The gene synteny of *clb* islands was plotted using clinker (v0.0.30) (*71*). Gene maps of *clbS* contigs were annotated with Prokka (v1.14.6) (*72*) and visualized with IGV (v2.19.7) (*73*).

The honeybee-related bacterial genomes were collected from NCBI as of August 2023. All genomes were quality-checked using CheckM (v1.2.2; lineage_wf mode) (*74*), and genomes with completeness ≤ 90% or contamination ≥ 5% were excluded. Protein-coding genes were predicted using Prodigal (*75*). ClbS homologs were detected using local protein-protein BLAST (blastp, v2.6.0) (*76*), with an e-value cutoff of 1e-5 and manual curation. The verified, functional ClbS-like sequences from Silpe *et al.*, 2022 (*29*) were used as reference sequences. The positive sequences were aligned with MAFFT (v7.505) (*77*) and manually curated. The phylogenetic tree was inferred using IQ-TREE 2 (v2.3.0) (*78*) and visualized using iTOL (https://itol.embl.de/) (*79*). Amino acid sequence identities of bee gut ClbS homologs to *E. coli* ClbS (APD28361.1) was evaluated using NCBI blastp webserver with default parameters.

Prophage-like structures were predicted according to a SOP for viral sequence identification (*80*) that combines VirSorter2 (v2.2.4) (*81*), CheckV (v1.0.3; database v1.5) (*82*), and DRAMv (v1.5.0) (*83*). The prophage-like islands were annotated and visualized using Pharokka (v1.7.3) (*84*).

### Statistical analyses and data visualization

Statistical analyses and data visualization were performed in R (v4.4.2) (*85*) and R studio (v2024.09.1) (*86*) or in Graphpad Prism (v10.3.1). Normal distribution and homogeneity of variances were assessed with Shapiro-Wilk test in base R and Levene’s test in R package “car” (v3.1.3) (*87*), respectively. R packages, including “tidyverse” (v2.0.0) (*88*), “beeswarm” (v0.4.0) (*89*), and “gghalves” (v0.1.4) (*90*), were used for data visualization.

Bee survival data were analyzed and plotted to Kaplan-Meier survival curves using R packages “survminer” (v0.5.0) (*91*) and “survival” (3.8.3) (*92*). In the pilot survival experiment, *p*-values were calculated using a pairwise log-rank test with Benjamini–Hochberg correction in R package “survminer”. For survival curves that combined two or three independent experiments, survival data were analyzed using a mixed-effect Cox proportional hazard model in R package “coxme” (v2.2.22) (*93*), with treatment groups as fixed effects and independent experiments as random effects. Assumptions of normality and homogeneity of variances were assessed and met before assessing the significance of fixed effects with ANOVA in R package “car”. Post-hoc pairwise comparisons between groups were conducted using estimated marginal means in R package “emmeans” (v1.10.6) (*94*), with Tukey-adjustment of the *p*-values.

Statistical significance of CFUs and fluorescence intensities was assessed using a Kruskal-Wallis test in base R, followed by a Dunn’s test in R package “FSA” (v 0.10.0) (*95*) with Benjamini–Hochberg correction for post-hoc pairwise comparisons. A Wilcoxon rank sum test with Benjamini–Hochberg correction in base R was used for comparing CFUs of two groups (*F. perrara* WT and *ΔclbB*). Linear regression was performed in base R.

## Acknowledgements

We thank Phillip Engel for kindly providing the *F. perrara ΔclbB* strain. We thank Erick V. S. Motta, Sean P. Leonard, and Gerald P. Maeda for their constructive suggestions, Kim Hammond for maintenance of bee hives and for assistance with figures, and Anna Webb and Richard Salinas for their help with fluorescence microscopy and flow cytometry, performed at the Center for Biomedical Research Support Microscopy and Flow Cytometry Facility at UT Austin (RRID:SCR_021756). This study was funded by the U.S. National Institutes of Health (R35GM131738 and R35GM158145) and the U.S. Army Research Office (W911NF-20-1-0195). E.P.B. is a Howard Hughes Medical Institute Investigator.

## Authors contributions

Y.S., N.A.M. conceived the study; Y.S., J.E.P., J.W.H.W., Y.L., M.A.A.R., E.P.B., N.A.M. designed experiments; Y.S., J.E.P., J.W.H.W., Y.L., M.A.A.R., T.D.J., J.M. performed experiments; N.A.M., E.P.B. provided resources and supervision; Y.S., N.A.M. wrote an original draft of the ms.; J.E.P., J.W.H.W., Y.L., M.A.A.R., E.P.B. also contributed to writing and editing. P.J.L. contributed the gene knockout method for *S. alvi* and gave comments on the ms.

## Supplementary Text

### The colibactin locus is ubiquitous in *F. perrara* genomes

So far there are only two sequenced isolates of *F. perrara*, strain PEB0191 (DSM 104328) from hives in Connecticut USA (*1*) and strain ESL0167 from Lausanne Switzerland (*2*). Both have the colibactin locus (Supplementary Fig. S1). There are also 9 *F. perrara* metagenome-assembled genomes (MAGs), which may not be complete, and 6 of these have the locus (Supplementary Fig. S1). The locus is absent in genomes of all other honeybee gut bacteria.

### *F. perrara* stimulates hindgut ROS to a similar level as other symbionts

Measurement of ROS in hindguts of gnotobiotic honeybees demonstrated that colonization by *F. perrara* increased ROS to a similar level as did three other bee gut symbionts, *S. alvi*, *G. apicola*, and *G. apis* (*p* > 0.38 between the symbionts), which were all higher than MD (*p* < 0.01) (Supplementary Fig. S4).

Elevated ROS generates the first line of pathogen defense and may explain the protective effect shortly after challenge (at day 1) from both WT and *ΔclbB* (Fig. 1D). However, the longer-term protection is mostly attributed to colibactin, as it is conferred by WT but not by *ΔclbB.* Bee gut symbionts modulate ROS production, which functions in determining host-symbiont specificity (*3*) and may also contribute to the observed protection against *S. marcescens* in previous studies (*4, 5*).

In mammalian systems, colibactin production appears to have complex effects on host immune responses. In murine models, *pks^+^ E. coli* did not alter intestinal inflammation under homeostatic conditions (*6*) but caused colibactin-dependent chronic colitis after a chemical disruption of mucosal integrity (*7*). On the other hand, the probiotic *E. coli* Nissle 1917 reduced intestinal inflammation of chemically induced colitis in a colibactin-dependent way (*8*). These studies suggest that effects of colibactin may vary depending on the producing strain and on gut homeostasis. Similarly, in *F. perrara,* colibactin may have multiple effects on host immune responses. Further study on the correlation between colibactin production by *F. perrara* and host immune responses will help to elucidate their potential interplay and combined effects on host health.

### Other attempts at investigating effects of *F. perrara* and colibactin on bee gut symbionts

We constructed pSL1-E2C-B2P*_recA_*eGFP, in which the promoter region of the *S. alvi recA* gene homolog was used for DNA damage reporting. However, it was not effective as MMC treatment failed to trigger a GFP signal (Supplementary Fig. S11). Three prophage-like islands were found in the genome of *S. alvi* wkB2 (Supplementary Fig. S12), but they were not induced by MMC or by *F. perrara* (Supplementary Fig. S13).

### *F. perrara* colonization has limited impact on the normal bee gut community

We inoculated a defined community of 8 core and 1 non-core (*Bartonella*) strains (Table S1) to gnotobiotic bees inoculated with *F. perrara* WT, *ΔclbB* or no *F. perrara* 5 days in advance (Supplementary Fig. S18A). Community structures in the hindguts of individual bees after another 5 days were estimated as relative and absolute abundances, using 16S rRNA gene amplicons classified into genera (Supplementary Fig. S18B and C).

*F. perrara ΔclbB* colonized less than WT, which is consistent with the prior CFU results. In general, absolute and relative abundances of taxa were similar across the three different *F. perrara* treatments. The exception was for *Gilliamella* which was negatively correlated with *Frischella,* whether WT or *ΔclbB* (*p* = 0.97), in linear regression analysis (Supplementary Fig. S18D). We reanalyzed community composition data of natural hive bees from two published studies (*9, 10*), and again found negative correlations between abundances of these two taxa (Supplementary Fig. S18E and F).

## Supplementary Figures

**Fig. S1.**
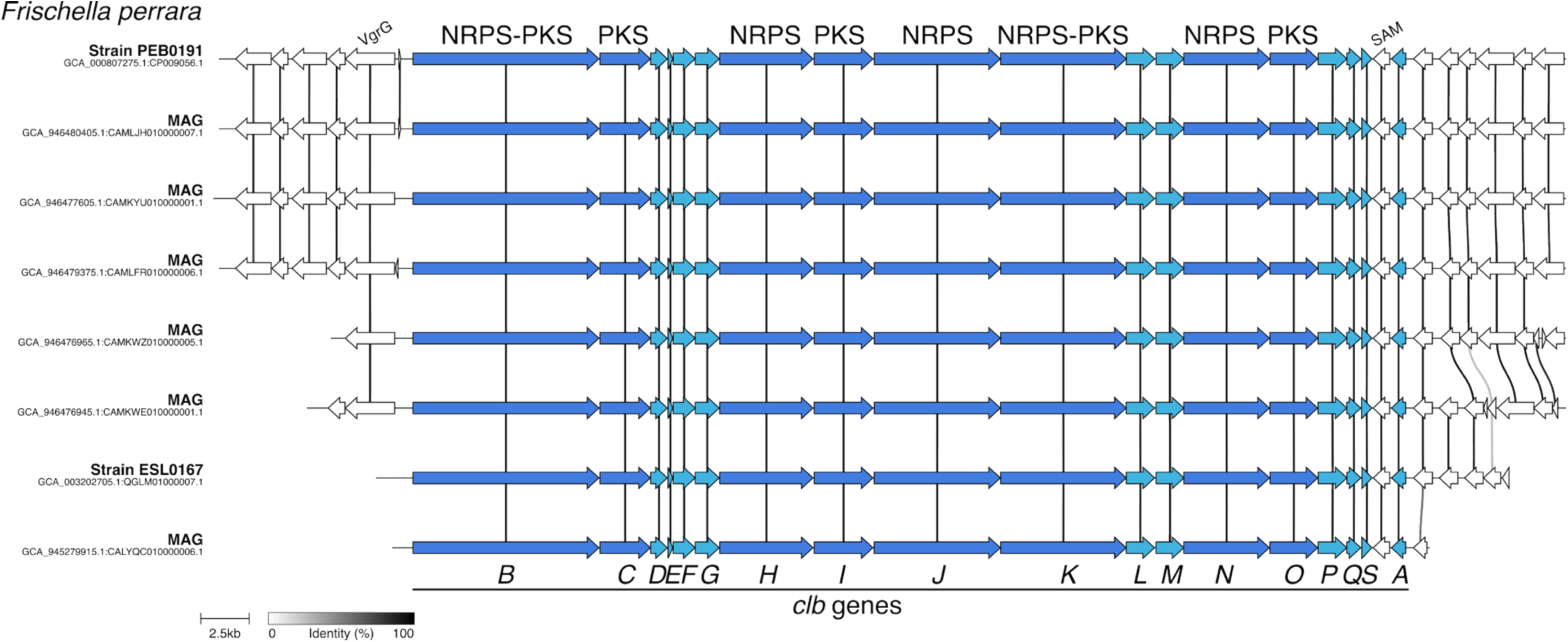
*Frischella perrara* genomes possess the highly conserved colibactin locus, with the exception of 3 metagenome-assembled genomes (MAGs: GCA_946699615.1, GCA_946477445.1, and GCA_025291255.1). Homologous genes are linked with lines, and line color depth indicates the percent amino acid identity as indicated by the key. Colibactin genes are in blue. Gene names (*clbA–S*) are given below. *F. perrara* (strain PEB0191) genes/proteins are detailed in the GenBank annotation (accession: NZ_CP009056.1). Abbreviations for neighboring genes: *VgrG*, type VI secretion system tip protein; SAM, radical *S*-adenosylmethionine protein; hypothetical proteins (white).

**Fig. S2.**
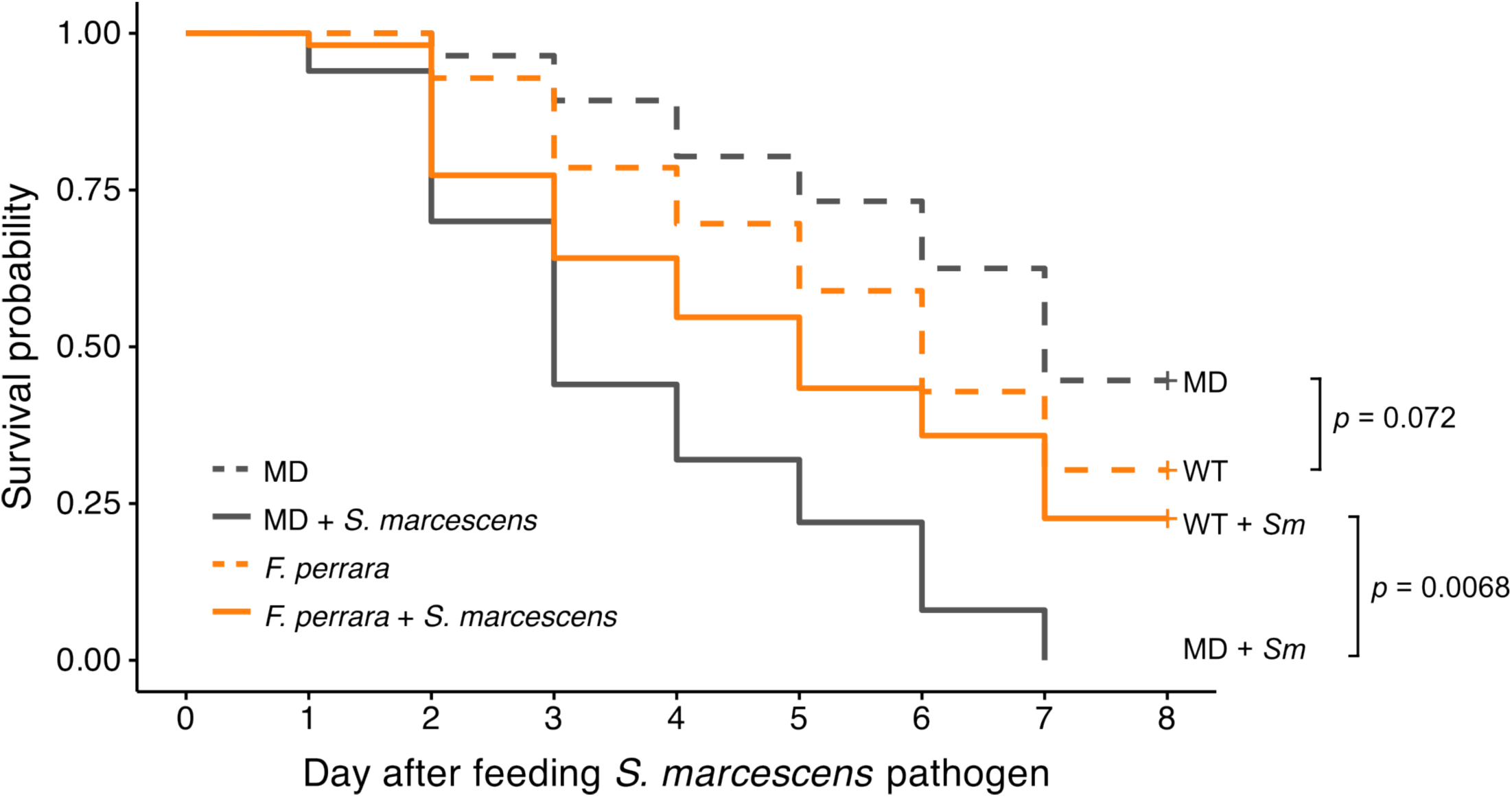
Survival of honeybees after pathogen challenge. Newly emerged bees initially received inoculation of either *Frischella perrara* symbiont or PBS control (microbiota-deprived, MD). The inoculation was conducted in 50 mL falcon tubes with breath holes. 150 µL cell suspension at OD_600_ of 1 in PBS were mixed with 150 µL sucrose water (*v/v*, 50:50) and added to tubes containing about 20 bees. Tubes were gently shaked to ensure the contact of bees with the mixture and kept statically for 5 min. The bees were subsequently moved to cup cages. After 5 days (day 0 in the plot), the bees were fed with the pathogen *Serratia marcescens* or PBS. Total *n* = 215 bees. *p*-values were calculated using a pairwise log-rank test with Benjamini–Hochberg correction.

**Fig. S3.**
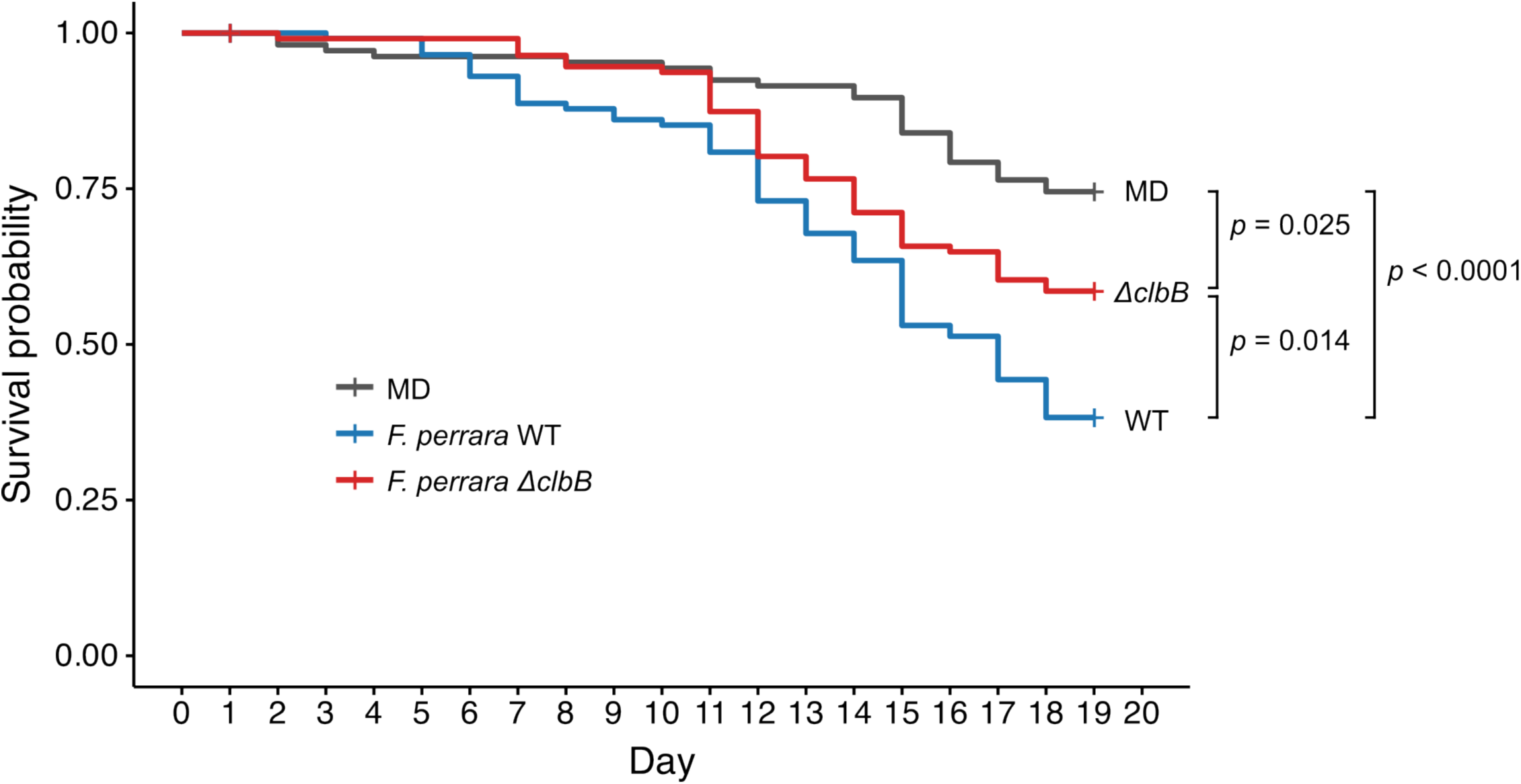
Survival of honeybees inoculated with *F. perrara* WT, *F. perrara ΔclbB* or not inoculated (MD). The Kaplan-Meier survival curves combined two independent experiments with bees sourced from different hives (∼20 bees per cup cage, 3 cages per group in each experiment). ANOVA test (*χ^2^* = 27.75, *df* = 2, *p* < 0.0001, *n* = 345) showed statistical significance among the treatment groups. Post-hoc pairwise comparisons were conducted using estimated marginal means.

**Fig. S4.**
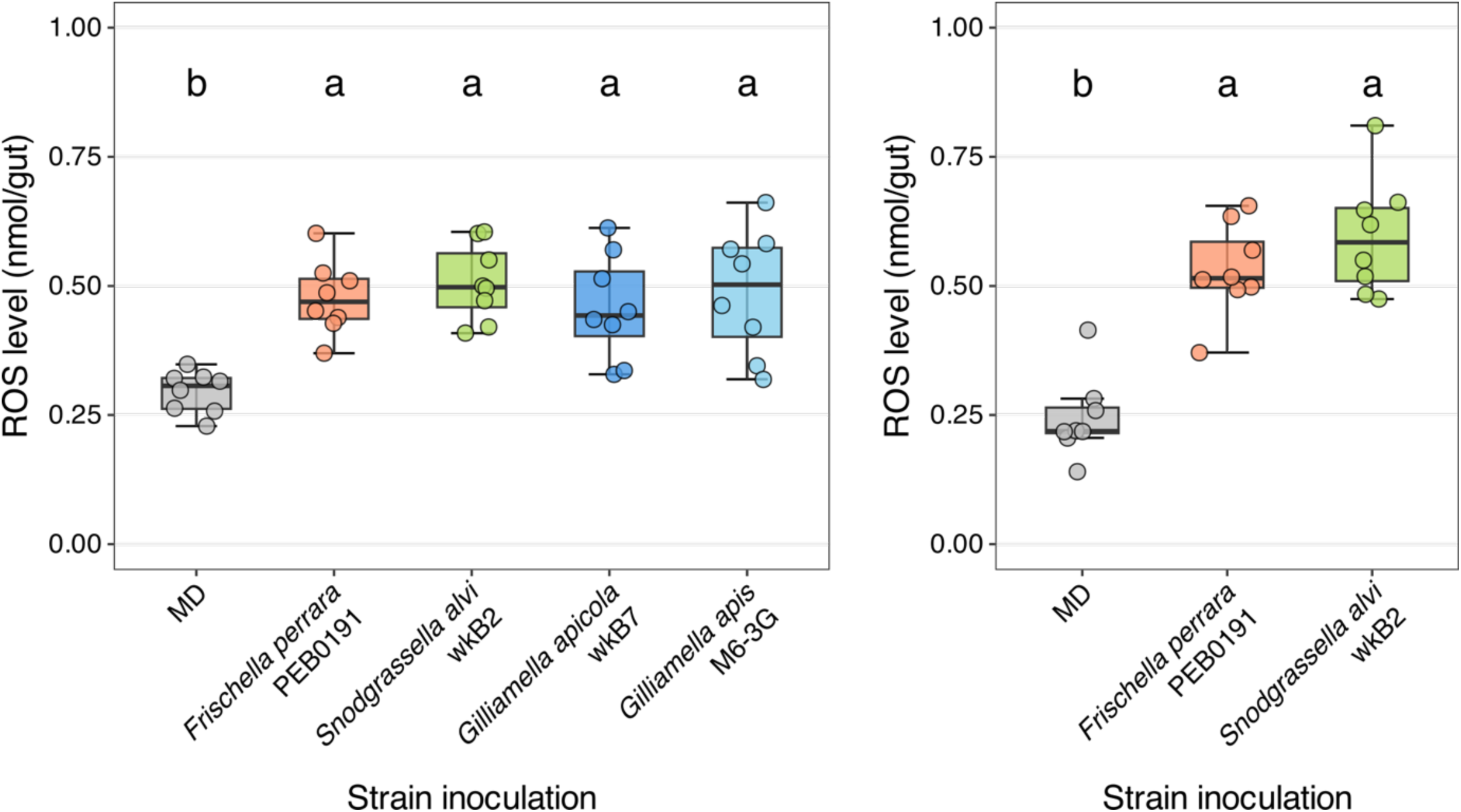
*F. perrara*, and other bee gut symbionts, increase ROS level in the hindgut of gnotobiotic honeybees. Results were from two independent experiments. ANOVA test (*F*(4, 35) = 7.903, *p* < 0.001, *n* = 40; *F*(2, 21) = 30.855, *p* < 0.0001, *n* = 24) showed statistical significance among groups in both experiments. Post-hoc pairwise comparisons were conducted using a Tukey’s pairwise comparisons test. Groups with different letters are significantly different (*α* = 0.05).

**Fig. S5.**
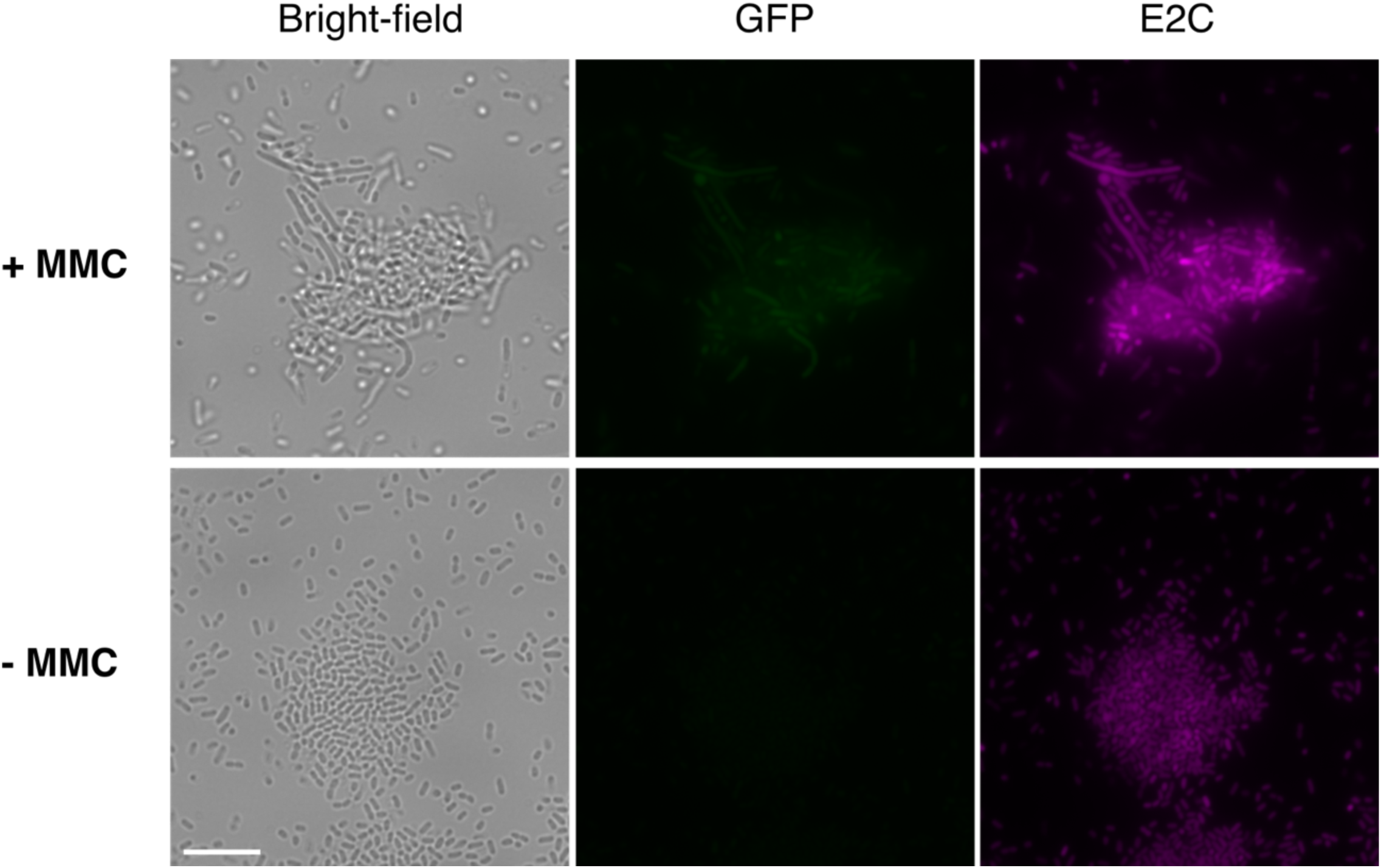
Mitomycin C (MMC, 0.05 µg/mL) treatment for 5 hours induced GFP in *S. marcescence* carrying pSL1-E2C-N10P*_recA_*eGFP. The elongated morphotype in the treated group is probably caused by arrested cell cycle. Scale bar = 10 µm.

**Fig. S6.**
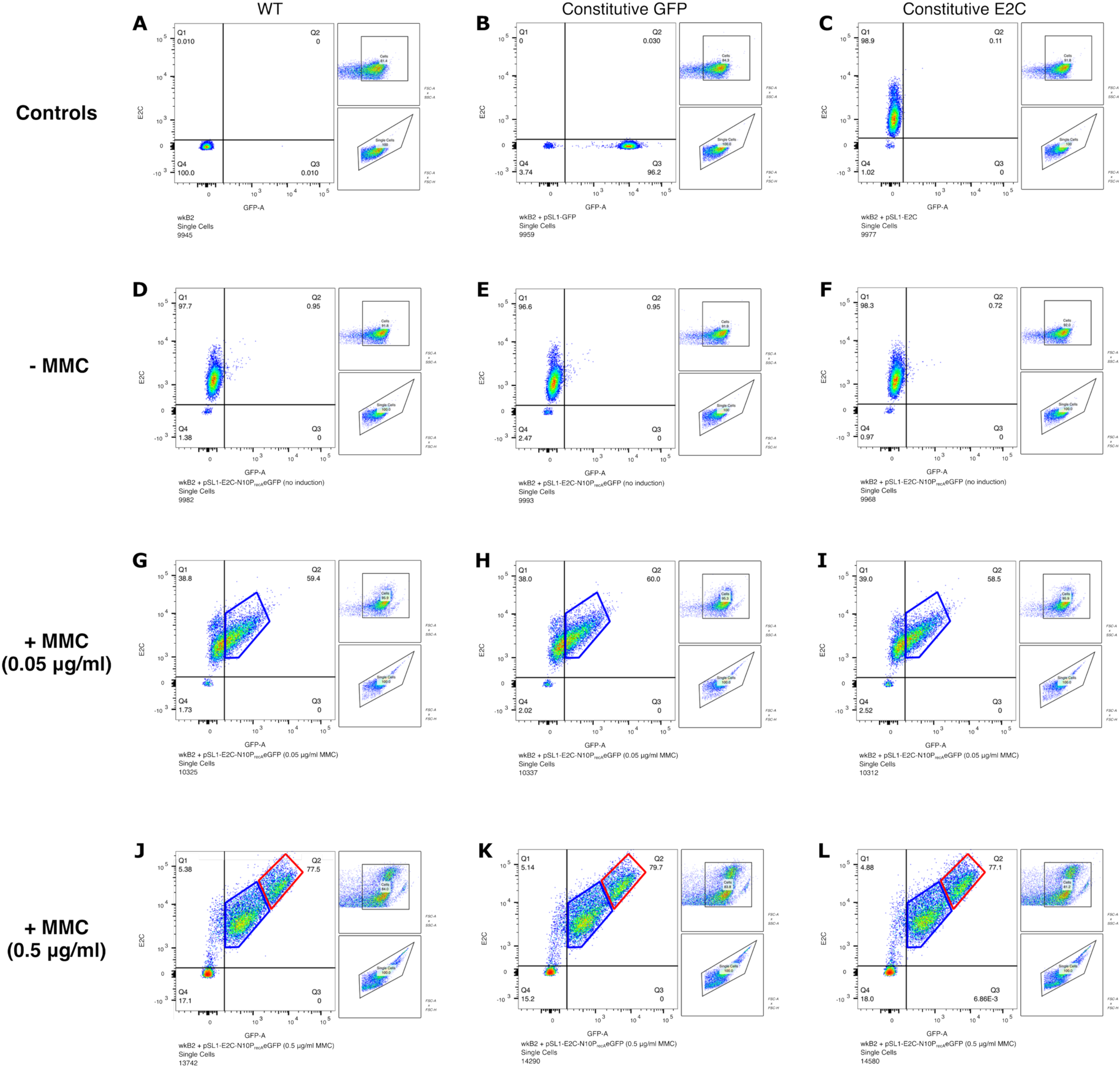
Fluorescence flow cytometry analysis of *S. marcescens* carrying pSL1-E2C-N10P*_recA_*eGFP, with or without MMC treatment. The left chart of each panel shows the intensity of GFP (x axis) and E2C (y axis), and right charts show the ancestry gating steps. The 4 quarters (Q1-4) were divided based on fluorescent and non-fluorescent controls (A-C). Q2 is positive in both GFP and E2C. *S. marcescens* plus pSL1-E2C-N10P*_recA_*eGFP was treated with no MMC (D-F), low MMC (G-I), or high MMC (J-L), three replicates each. Both low and high concentrations of MMC triggered a population with increased GFP intensity (blue gated). High concentration triggered another population with both GFP and E2C intensities increased (red gated) and also more cells that lost both fluorescence in Q4.

**Fig. S7.**
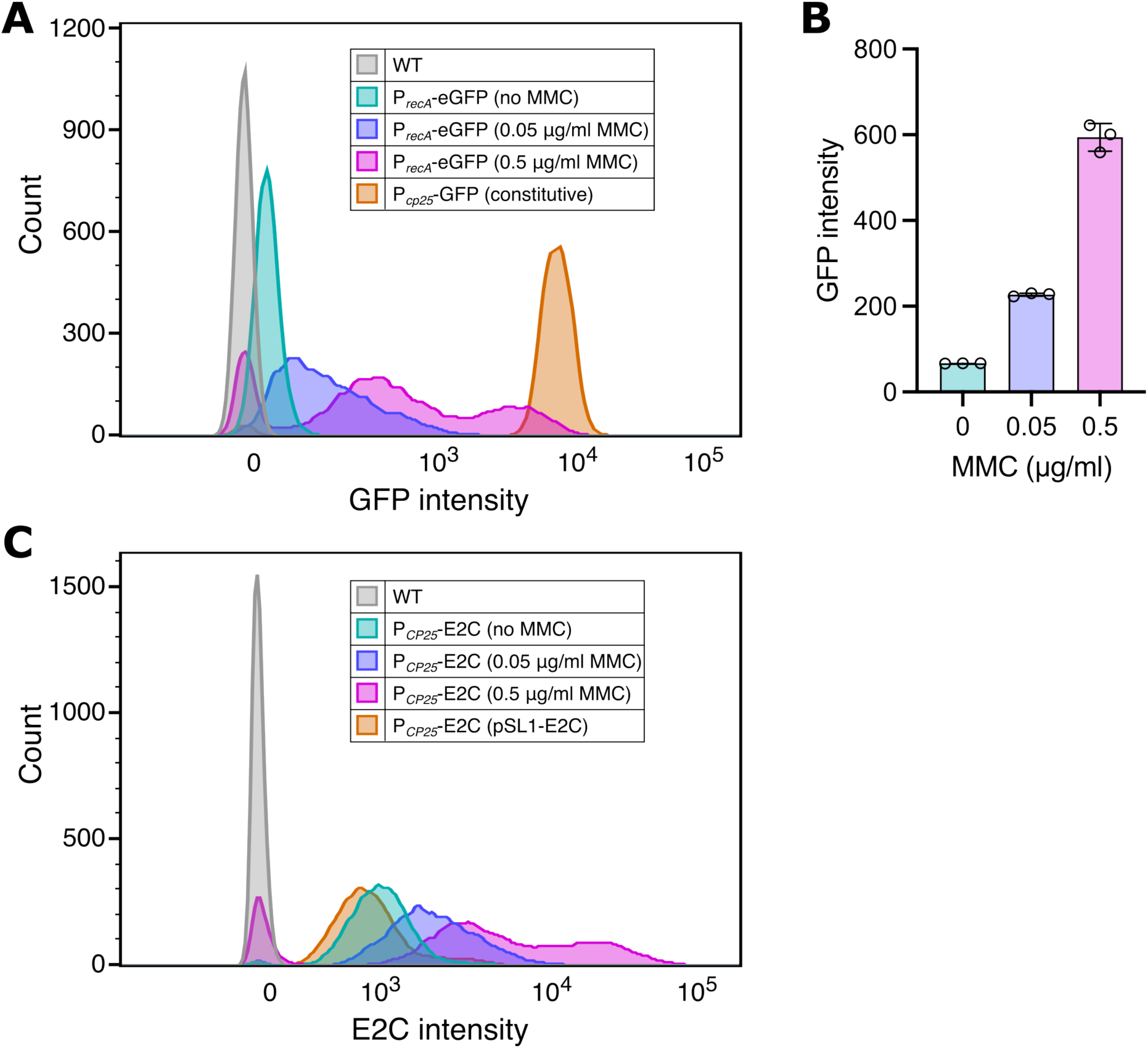
Fluorescence intensities of *S. marcescens* carrying pSL1-E2C-N10P*_recA_*eGFP, with or without MMC treatment. (A) GFP intensity under different MMC concentrations. WT and WT + pSL1-GFP (constitutive GFP) were used as controls. (B) Median GFP intensity (± SD) increased with MMC concentration. (C) E2C intensity under different MMC concentrations. WT and WT + pSL1-E2C were used as controls. All samples (except WT) constitutively express E2C.

**Fig. S8.**
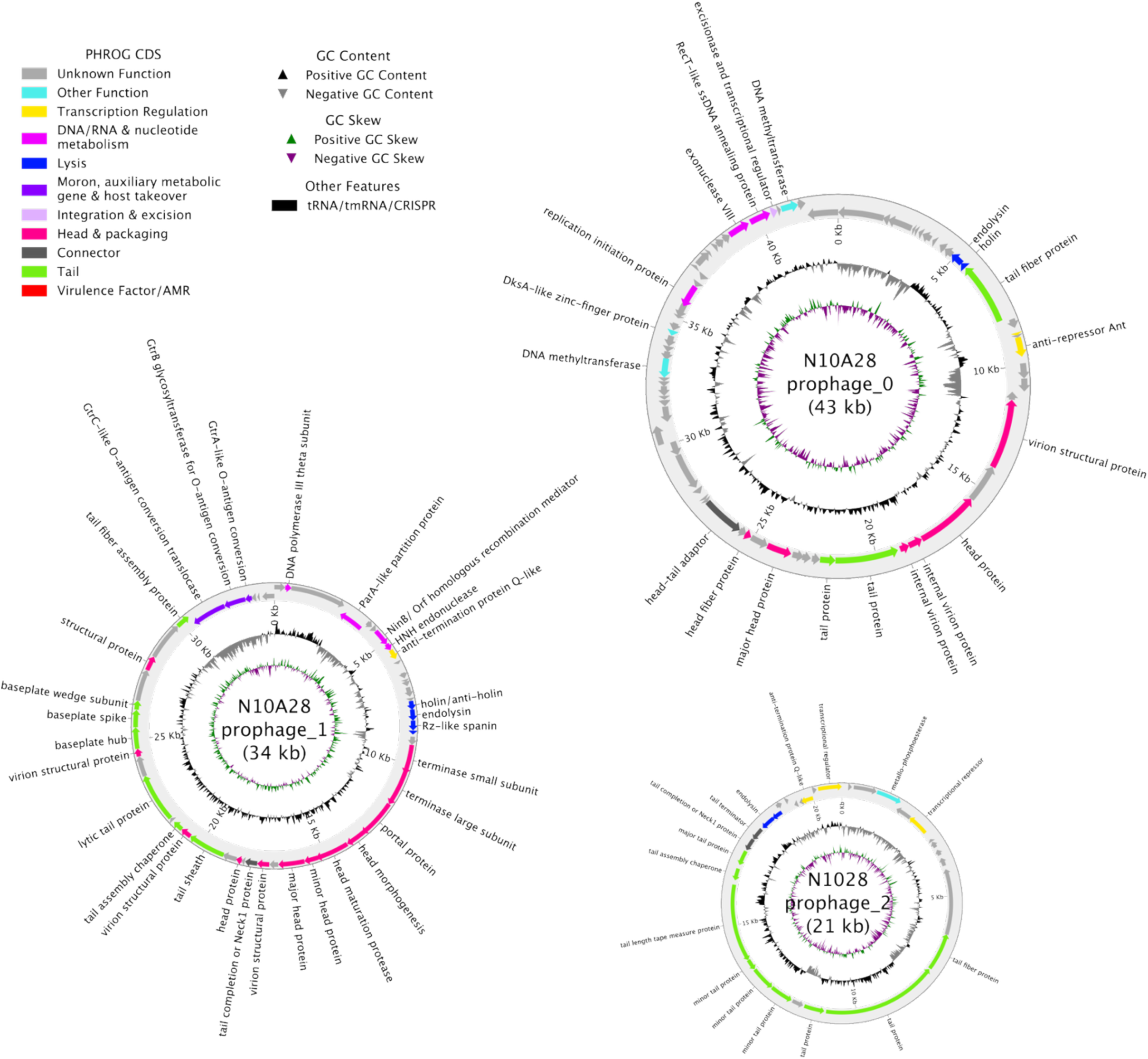
Three predicted prophage-like structures in the *Serratia marcescens* N10A28 genome (GenBank: CP033623.1). Genomic coordinates: 1,400,308–1,434,310 (Prophage_1); 2,928,641–2,949,521 (Prophage_2); 3,342,031–3,385,057 (Prophage_0).

**Fig. S9.**
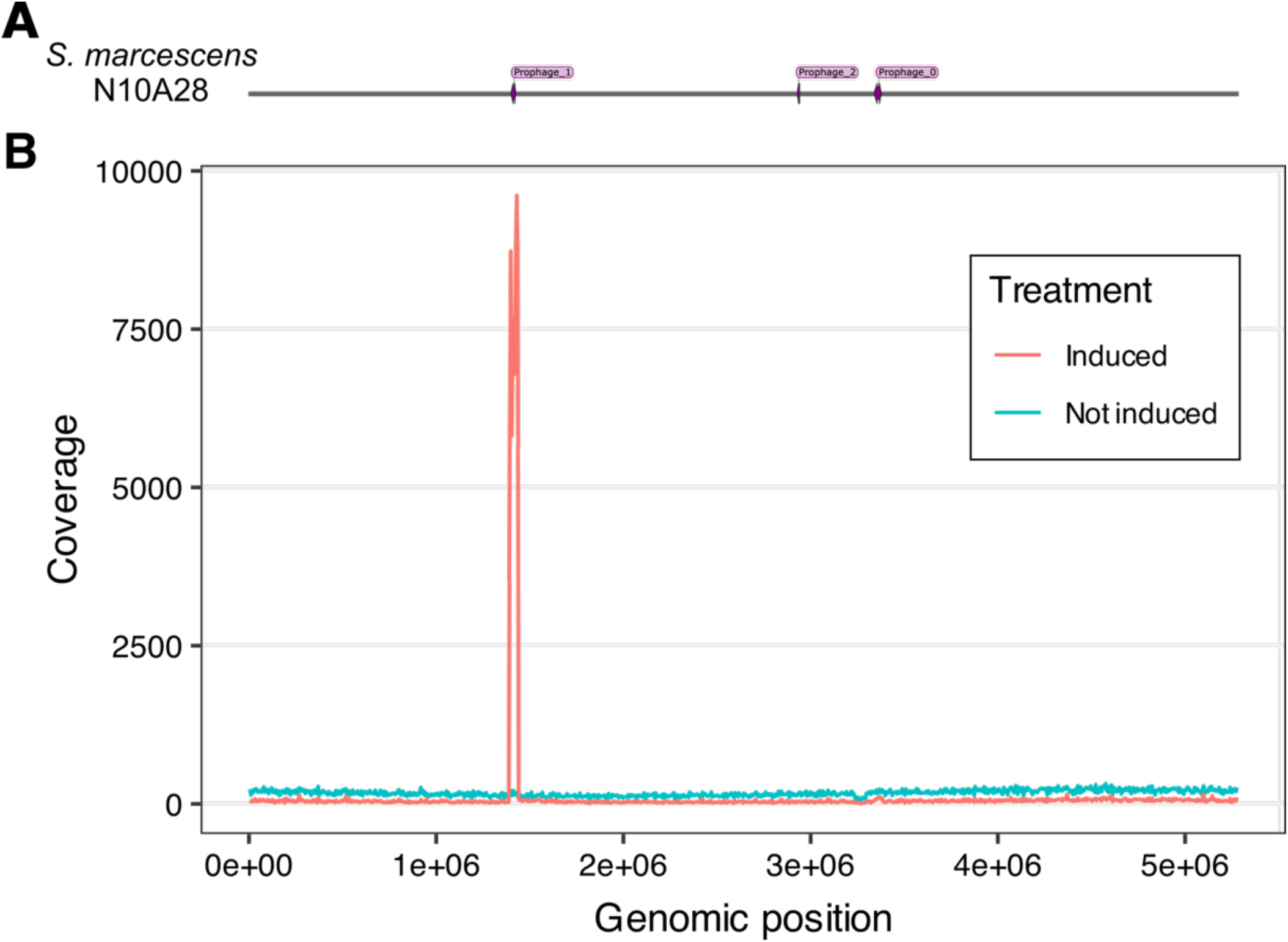
MMC induced a prophage region in *S. marcescens* N10A28. (A) Purple loci showing positions of the three prophage-like structures in the genome. (B) MMC (0.5 µg/mL) treatment for 5 hours induced lytic replication of Prophage_1; the induced genomic region (1,390,922–1,435,747) encompasses the predicted region (1,400,308–1,434,310). The other two prophage-like regions were not induced. Without MMC, all prophage-like structures were not induced.

**Fig. S10.**
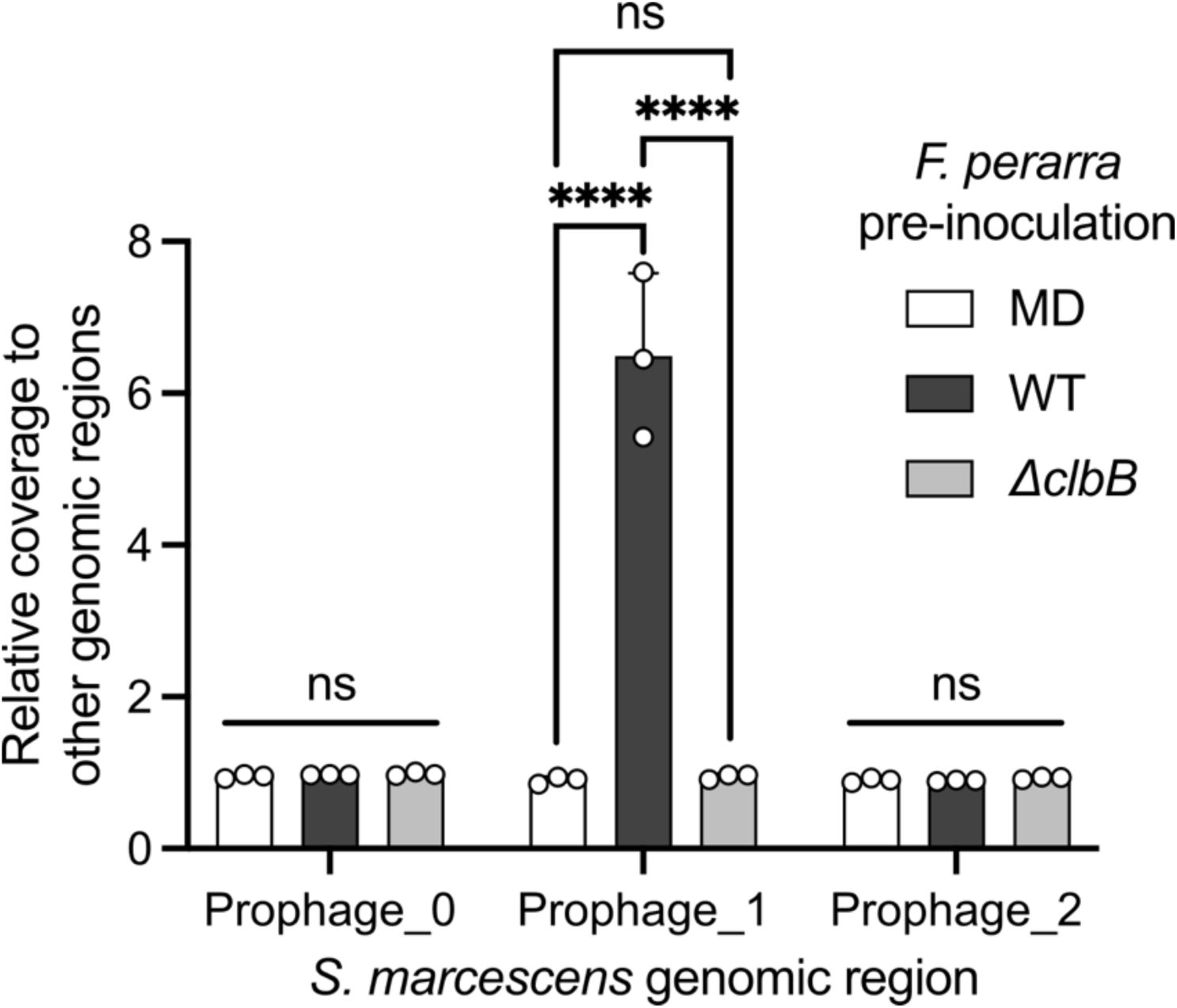
Relative coverage of *S. marcescens* prophage-like regions to other genomic regions in bee guts pre-colonized with *F. perrara* WT, *ΔclbB* or MD. Prophage_1 was induced by *F. perrara* WT but not by *F. perrara ΔclbB*. The other two prophage-like structures were not induced under all tested conditions. Statistical differences were assessed using a one-way ANOVA with a Tukey’s multiple comparisons test (Graphpad Prism). Statistical significance: ****, *p* < 0.0001; ns, not significant.

**Fig. S11.**
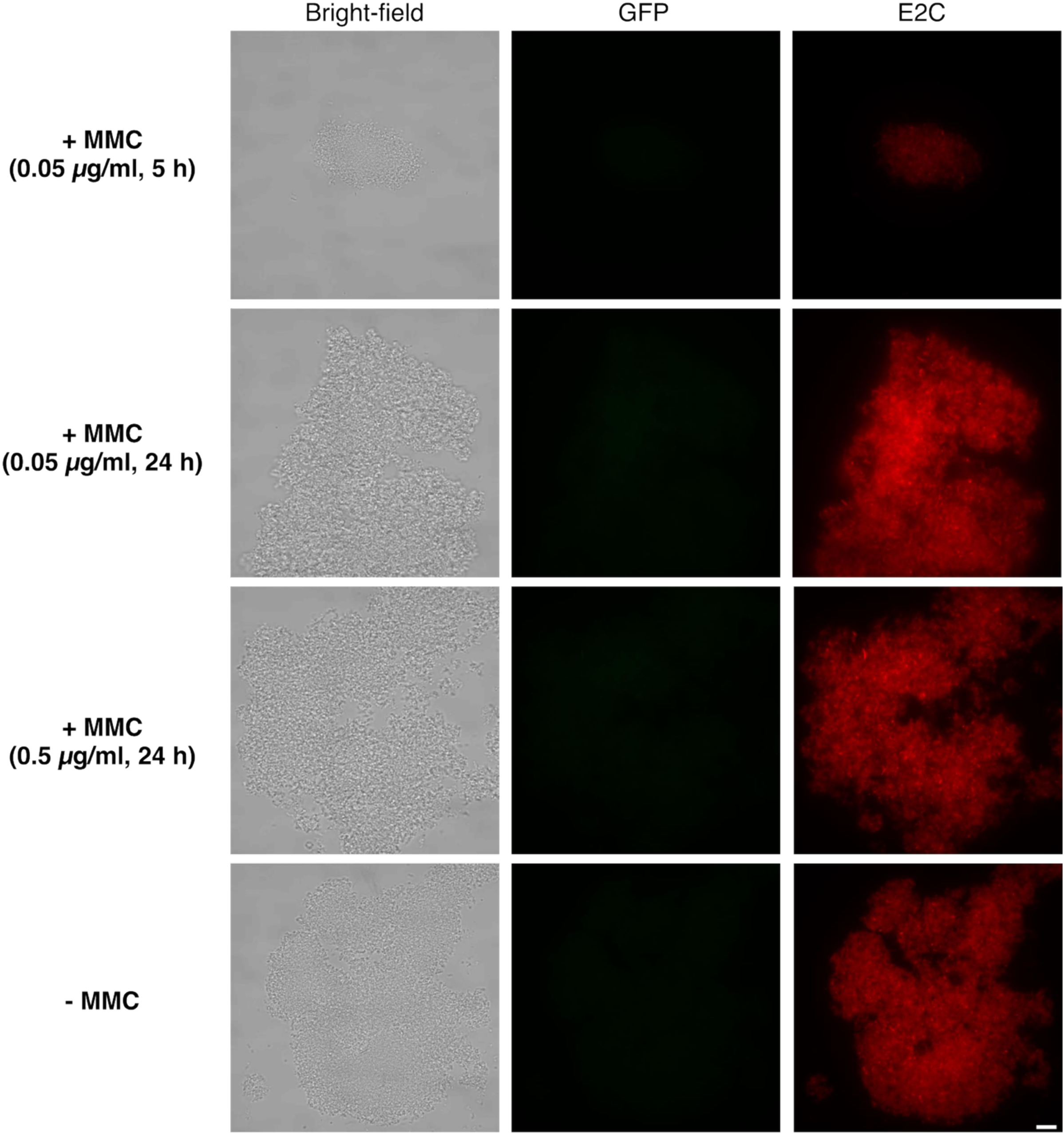
MMC treatment failed to induce GFP signal in *S. alvi* wkB2 carrying pSL1-E2C-B2P*_recA_*eGFP. Scale bar = 10 µm.

**Fig. S12.**
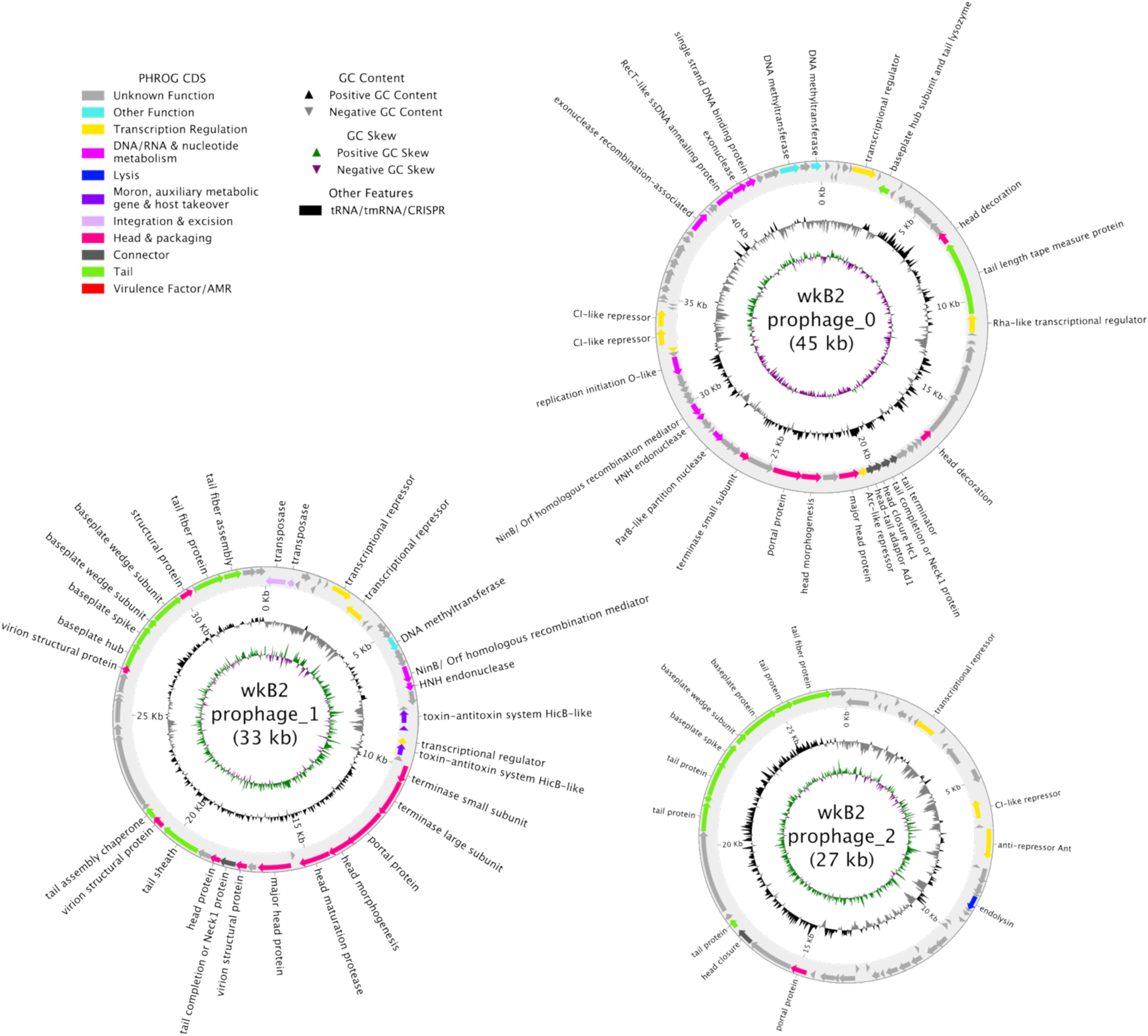
Predicted prophage-like structures in the *Snodgrassella alvi* wkB2 genome (CP007446.1). Genomic coordinates: 547,445–580,557 (Prophage_1); 1,033,694–1,060,775 (Prophage_2); 2,046,103–2,091,026 (Prophage_0).

**Fig. S13.**
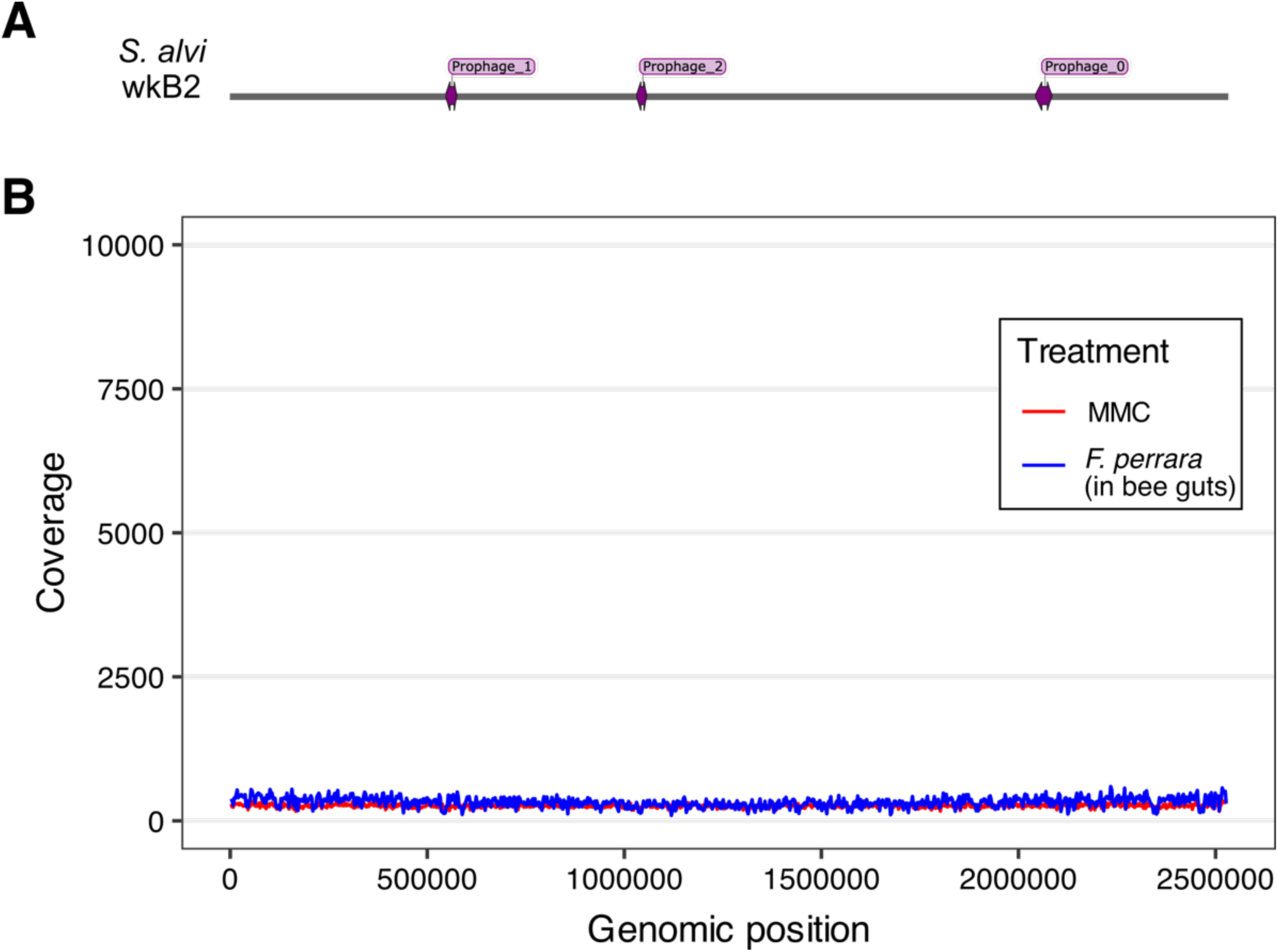
MMC and *F. perrara* failed to induce lytic replication of prophage-like regions in *S. alvi* wkB2. (A) Purple loci show positions of the three prophage-like structures in the genome. (B) MMC (0.5 µg/mL) treatment for 5 hours or *F. perrara* pre-inoculation did not show prophage induction in wkB2. Bees were pre-inoculated with *F. perrara* WT, after 6 days, inoculated with *S. alvi ΔclbS*, and hindguts were collected after another 4 days.

**Fig. S14.**
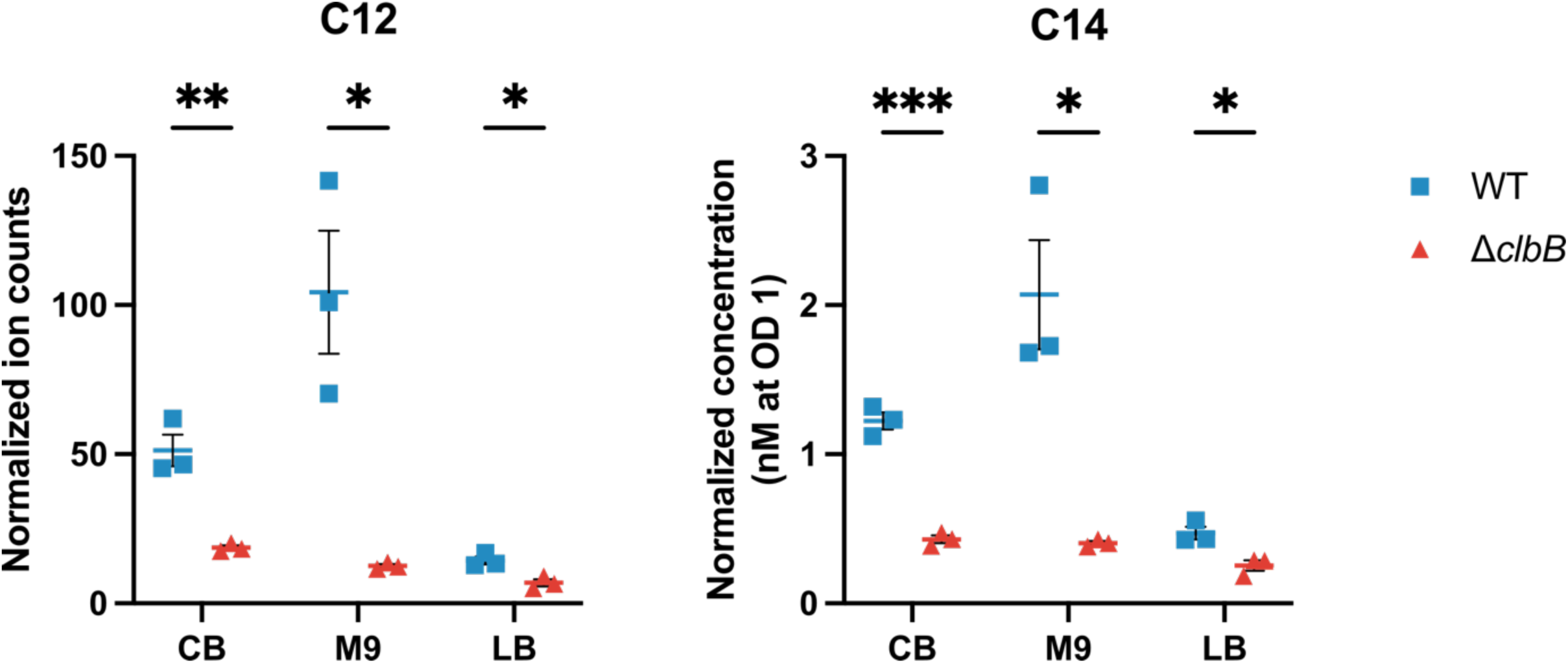
LC–MS measurement of *N*-lauryl-D-Asn (C12 prodrug) and *N*-myristoyl-D-Asn (C14 prodrug) in cell pellets of *F. perrara* WT and *ΔclbB* cultured with different media. Data are mean ± SEM; *n* = 3 biological replicates. Statistical differences between WT and *ΔclbB* were assessed using a two-tailed unpaired t-test (Graphpad Prism). Statistical significance: ***, *p* < 0.001; **, *p* < 0.01; *, *p* < 0.05. For the M9 groups, strains were cultured in CB first and cell pellets were recollected, washed, and resuspended in M9 medium. *F. perrara* showed very weak growth in LB medium (final OD ∼1/10 of that in CB).

**Fig. S15.**
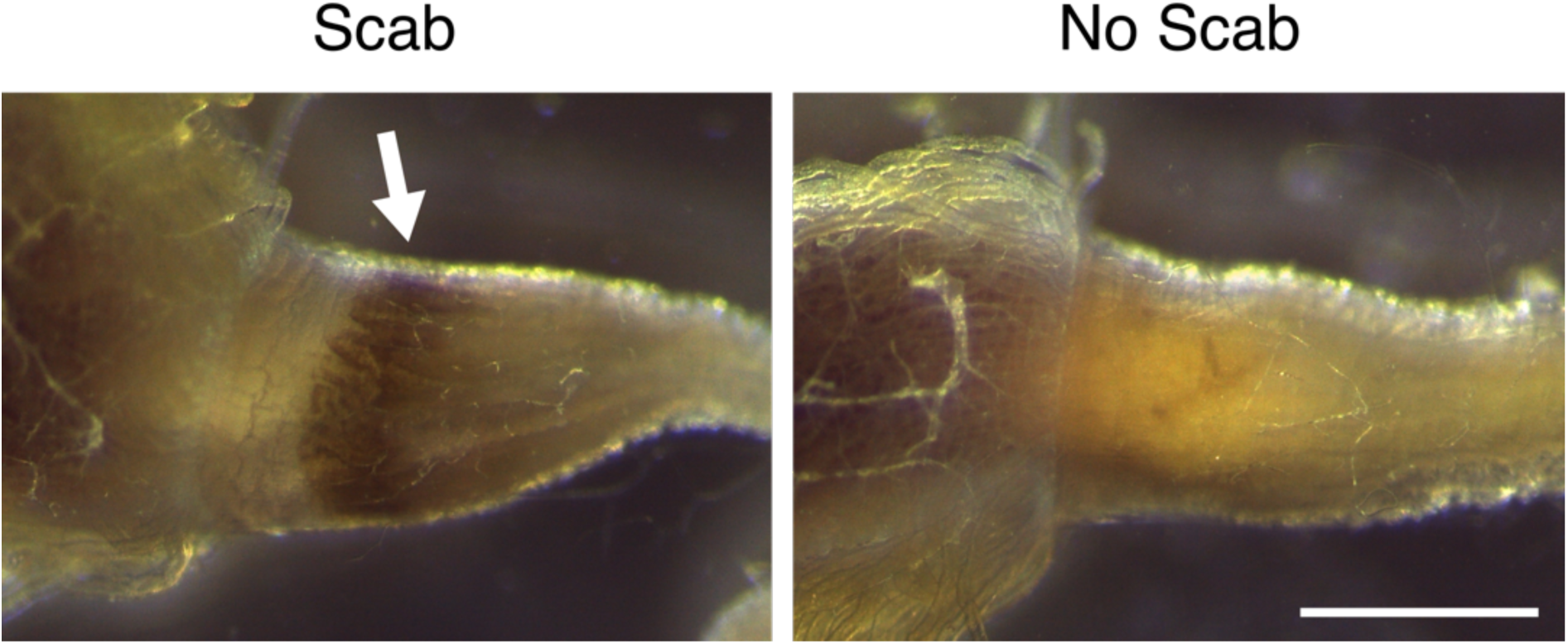
Scab-like phenotype in the pylorus of the bee gut caused by *F. perrara* colonization. Arrow indicates the melanized region. Scale bar = 0.5 mm.

**Fig. S16.**
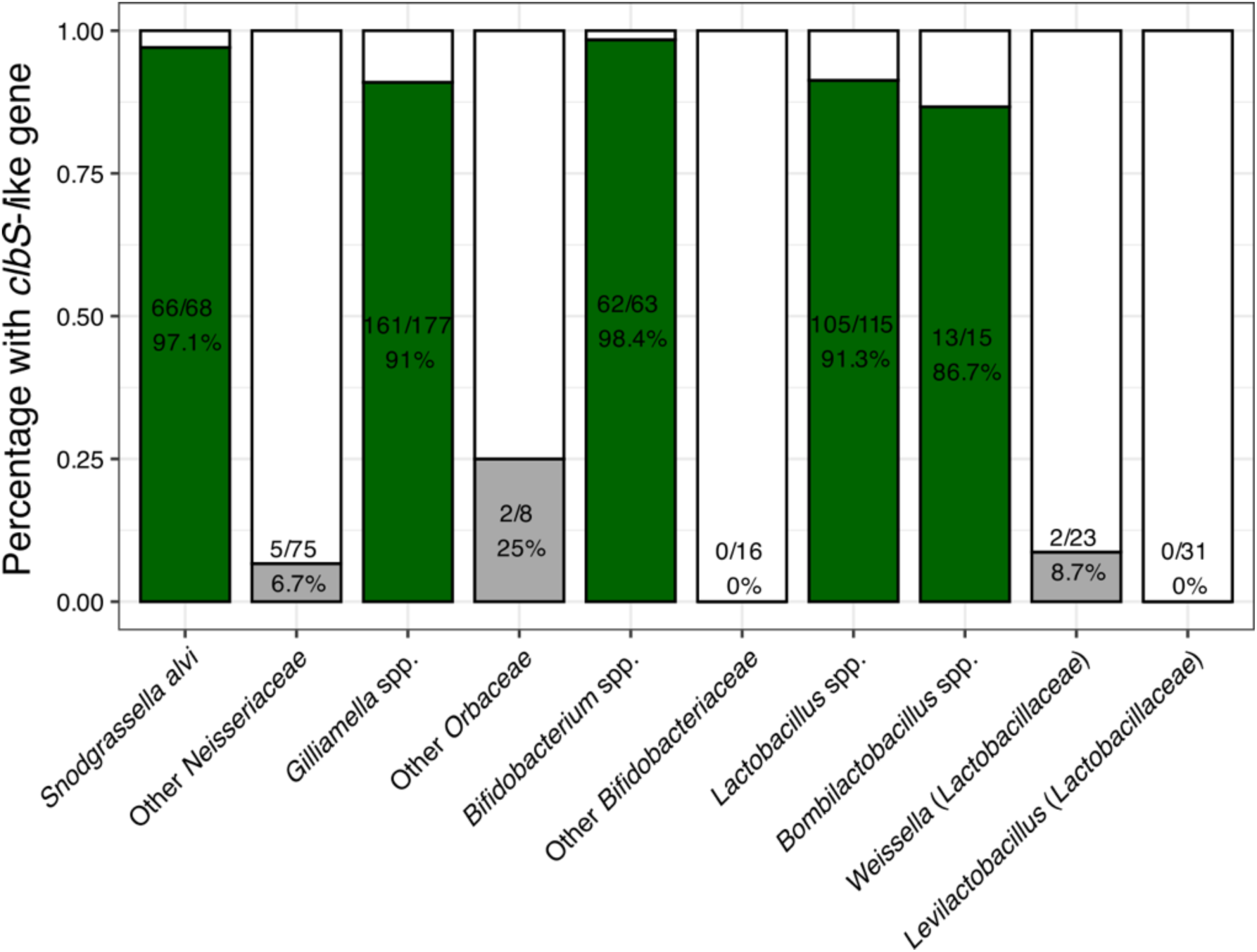
Presence of *clbS*-like gene in genomes of bee gut core taxa (in green) and closely related taxa from other environments (in grey). Positive/total genome number and percent are given in the plot. Genomes and sequences are detailed in Supplementary Data S1.

**Fig. S17.**
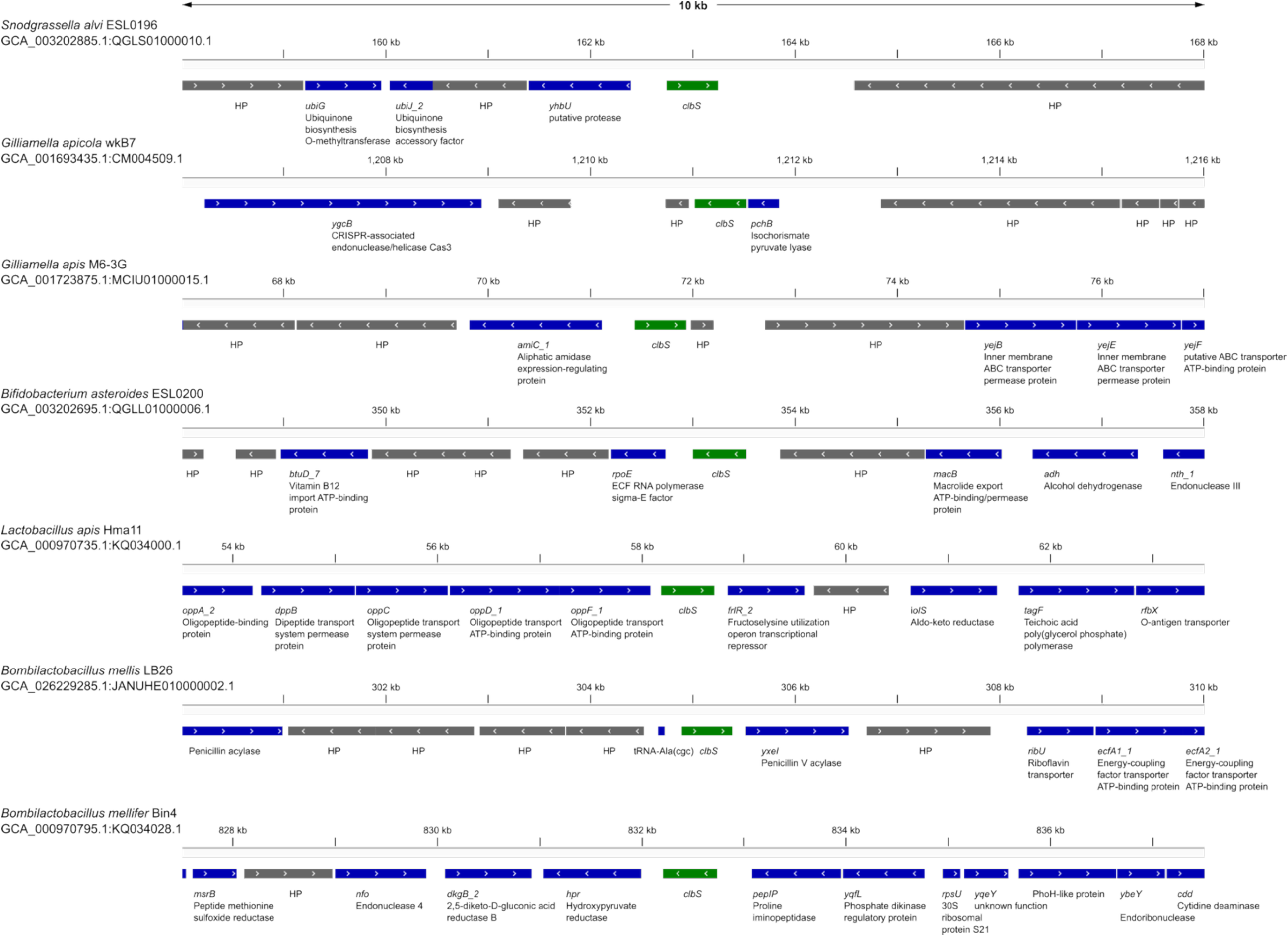
Gene maps of the tested *clbS* homologs in Fig. 3B. The *clbS* genes are in green. Neighboring genes with annotated functions are in blue (gene abbreviations and names are given), and those encoding hypothetical proteins (HPs) are in grey.

**Fig. S18.**
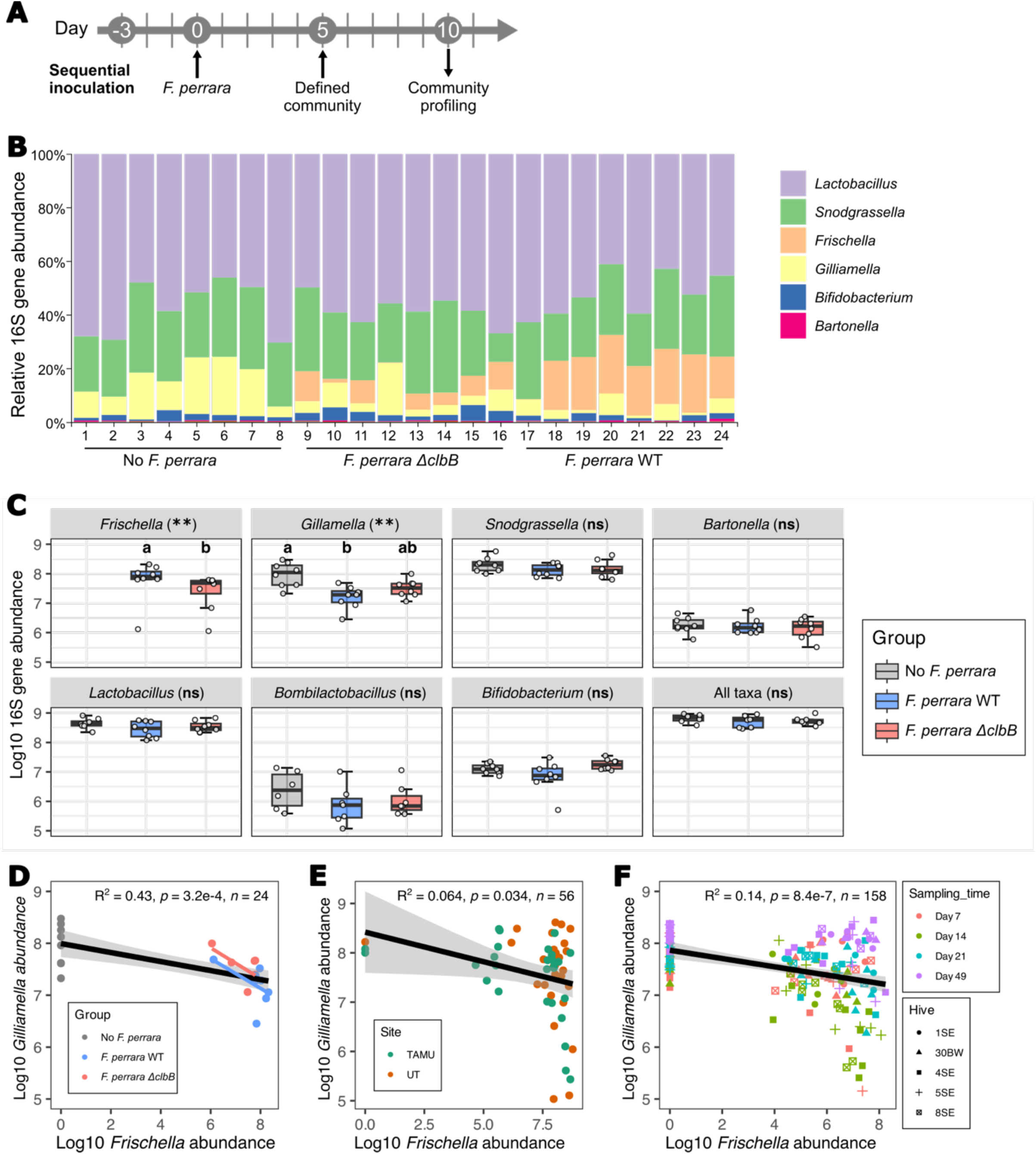
Levels of *Frischella* and *Gilliamella* are negatively correlated in honeybee guts, but *Frischella* presence does not otherwise affect community composition. (A) Timeline of the sequential inoculation of *F. perrara* and a defined community of honeybee gut bacterial symbionts. (B) Relative abundance of different taxa (classified into genus-level) in the defined community assay. (C) Absolute abundance of different taxa in the defined community assay. Wilcoxon rank sum test indicated a statistically significant difference between *F. perrara* WT and *ΔclbB* (*p* = 0.007). Kruskal–Wallis test showed no statistical significance in other taxa except *Gilliamella* (*χ^2^* = 9.695, *df* =2, *p* = 0.008, *n* = 24). Post-hoc pairwise comparisons between *Gilliamella* groups were conducted using Dunn’s test. Statistical significance: **, *p* < 0.01; ns, not significant. Groups with different letters are significantly different (*α* = 0.05). (D) *F. perrara* and *Gilliamella* spp. showed negative correlation in the defined community assay. (E) and (F) Negative correlation between *Frischella* and *Gilliamella* in the gut of hive sourced bees. Data from Powell *et al.*, 2021 and 2023 (*9, 10*).

## Supplementary Tables

**Table S1.** Bacterial strains used in this study.

| Species | Strain | Antibiotic Marker | Source |
| --- | --- | --- | --- |
| *Frischella perrara* | PEB0191 | - | (11) |
| *Frischella perrara* | PEB0191 $\Delta clbB$ | - | (12) |
| *Snodgrassella alvi* | wkB2<sup>a</sup> | - | (13) |
| *Snodgrassella alvi* | wkB2 $\Delta clbS$ | *ampR* | This study |
| *Serratia marcescens* | N10A28 | - | (10) |
| *Gilliamella apicola* | wkB7<sup>a</sup> | - | (13) |
| *Gilliamella apis* | M6-3G<sup>a</sup> | - | (14) |
| *Bifidobacterium asteroides* | wkB338<sup>a</sup> | - | (15) |
| *Lactobacillus helsingborgensis* | wkB8<sup>a</sup> | - | (16) |
| *Lactobacillus kullabergensis* | wkB10<sup>a</sup> | - | (16) |
| *Bombilactobacillus mellis* | Hon2N<sup>a</sup> | - | (17, 18) |
| *Bombilactobacillus mellifer* | Bin4N<sup>a</sup> | - | (17, 18) |
| *Bartonella apis* | PEB0150<sup>a</sup> | - | (19) |
| *Escherichia coli* | DH5α | - | NEB #C2987H |
| *Escherichia coli* | MFDpir | - | (20) |
| *Escherichia coli* | BW25113 | - | (21) |

<sup>a</sup> Strains included in the defined community.

**Table S2.** Plasmids used in this study.

| Plasmid | Function | Antibiotic Marker | Source |
| --- | --- | --- | --- |
| pSL1-GFP | Constitutive expression of GFP under CP25 promoter | Kan | (22) |
| pSL1-RCP | Constitutive expression of Red Chrome Protein under CP25 promoter | Kan | (22) |
| pSL1-E2C | Constitutive expression of E2-Crimson under CP25 promoter | Kan | (22) |
| pSL1-E2C-N10P<sub>recA</sub>eGFP | Constitutive expression of E2-Crimson under CP25 promoter and inducible expression of eGFP by DNA damage under *recA* promoter of *Serratia marcescens* N10A28 | Kan | This study |
| pSL1-E2C-B2P<sub>recA</sub>eGFP | Constitutive expression of E2-Crimson under CP25 promoter and inducible expression of eGFP by DNA damage under *recA* promoter of *Snodgrassella alvi* wkB2 | Kan | This study |
| pBeloBAC11 | Empty bacterial artificial chromosome (BAC) | Cm | NEB |
| BAC-*pks* | BAC with *pks* genomic island for colibactin production | Cm | (23) |
| P<sub>R</sub>-*lux* | Bioluminescent reporter for DNA damage | Kan | (24) |
| pTrcHis2_A | Empty pTrc vector for *clbS*-like genes cloning | Amp | Thermo Fisher Scientific |
| pTrc-*clbS* | Expression of WT *clbS* from *E. coli* CFT073 under Trc promoter | Amp | This study |
| pTrc-*clbS*<sub>Snod</sub> | Expression of *clbS*-like gene from *Snodgrassella alvi* ESL0196 | Amp | This study |
| pTrc-*clbS*<sub>Gillia1</sub> | Expression of *clbS*-like gene from *Gilliamella apicola* wkB7 | Amp | This study |
| pTrc-*clbS*<sub>Gillia2</sub> | Expression of *clbS*-like gene from *Gilliamella apis* M6-3G | Amp | This study |
| pTrc-*clbS*<sub>Bifido</sub> | Expression of *clbS*-like gene from *Bifidobacterium asteroides* ESL0200 | Amp | This study |
| pTrc-*clbS*<sub>Lacto</sub> | Expression of *clbS*-like gene from *Lactobacillus apis* Hma11 | Amp | This study |
| pTrc-*clbS*<sub>Bombi1</sub> | Expression of *clbS*-like gene from *Bombilactobacillus mellis* LB26 | Amp | This study |
| pTrc-*clbS*<sub>Bombi2</sub> | Expression of *clbS*-like gene from *Bombilactobacillus mellifer* Bin4 | Amp | This study |

**Table S3.**
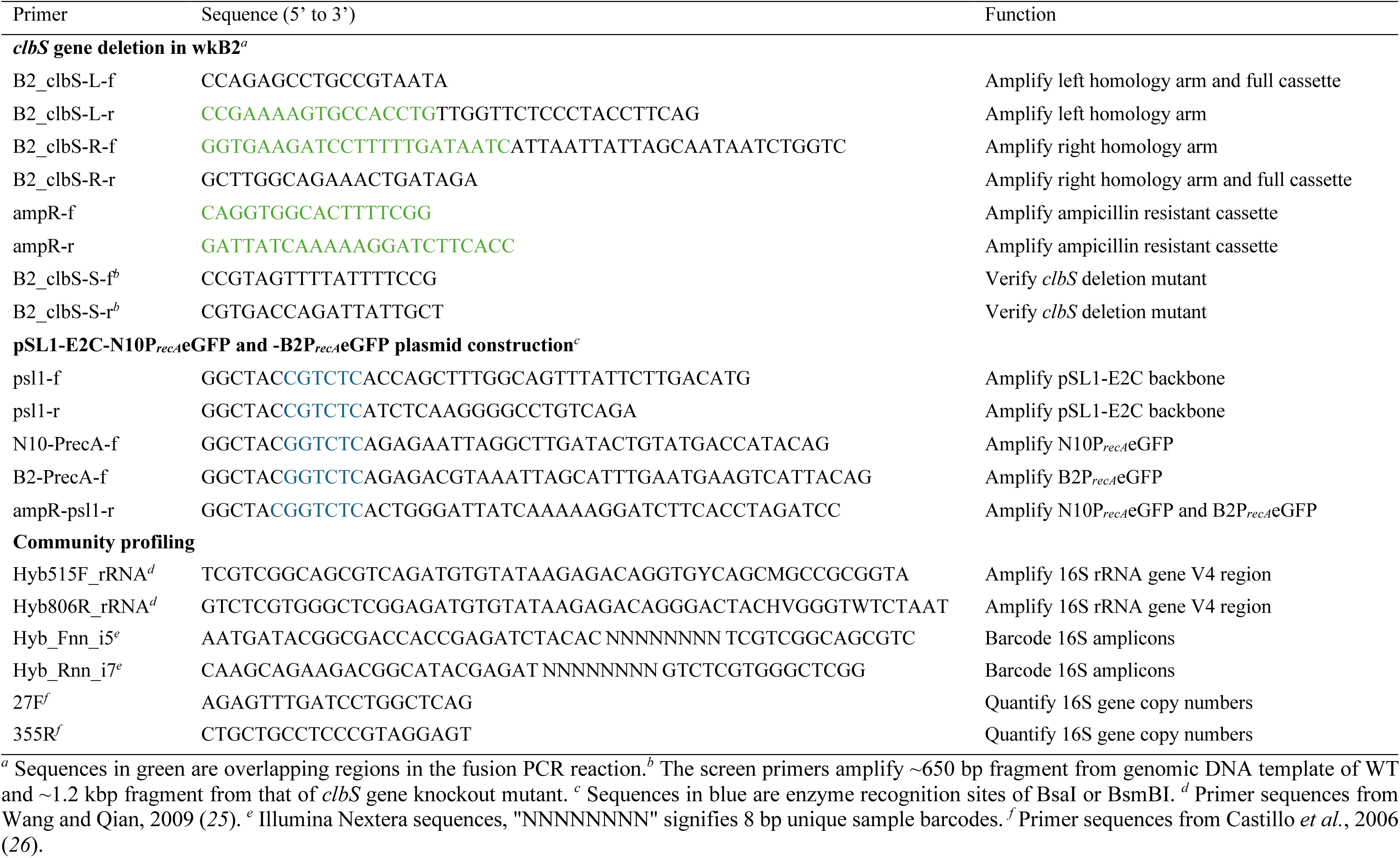
Primers used in this study.

**Table S4.**
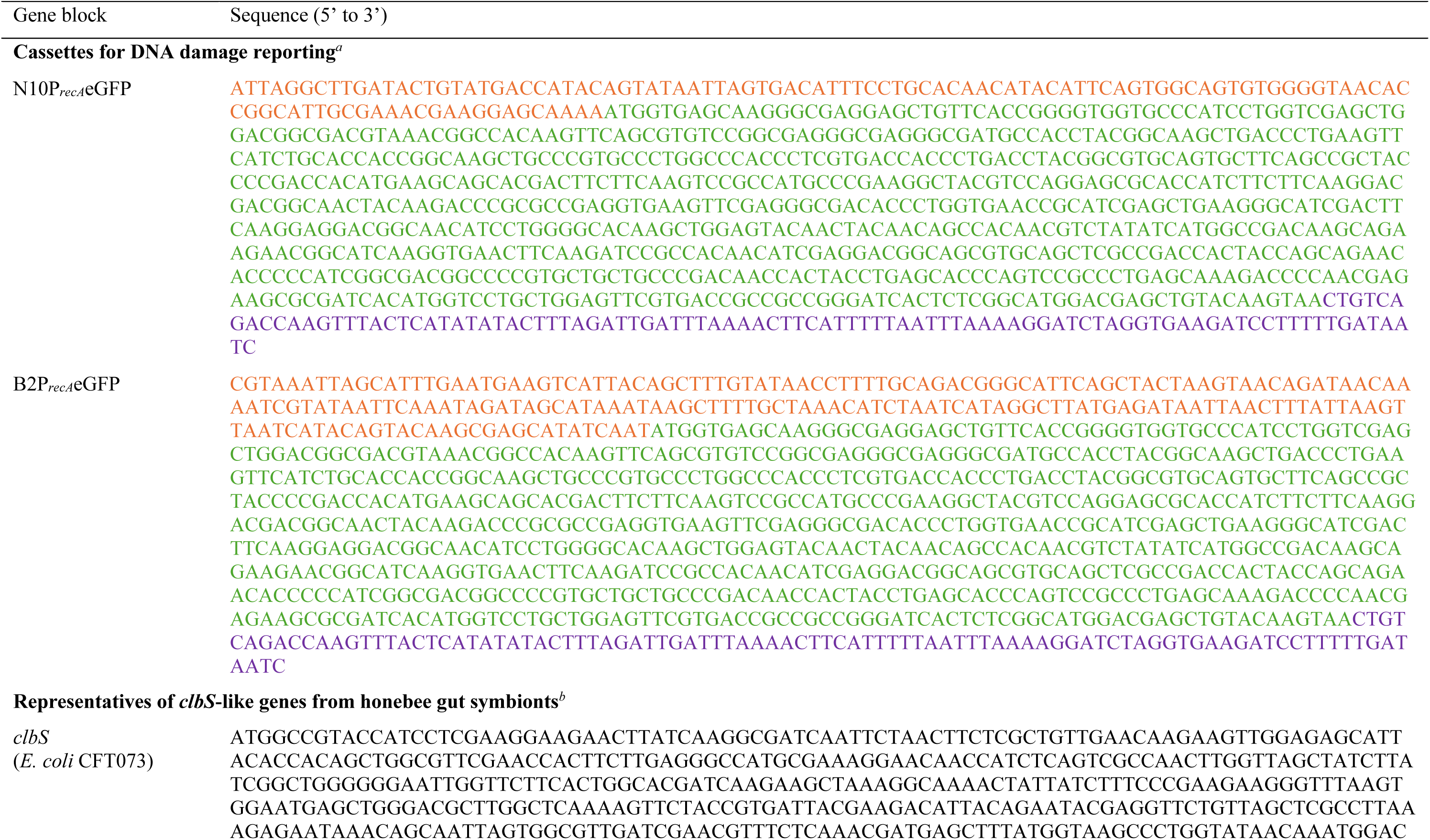
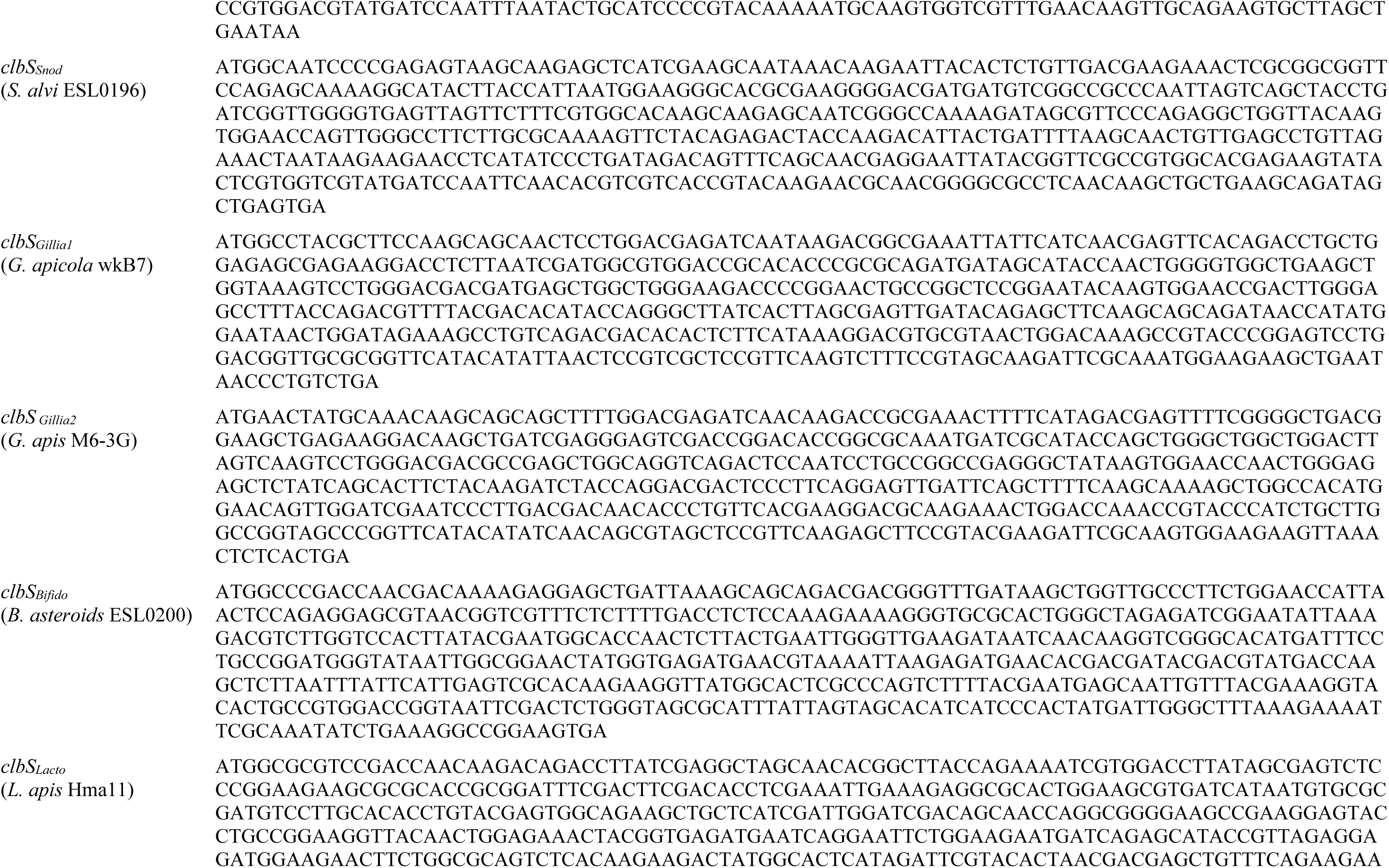
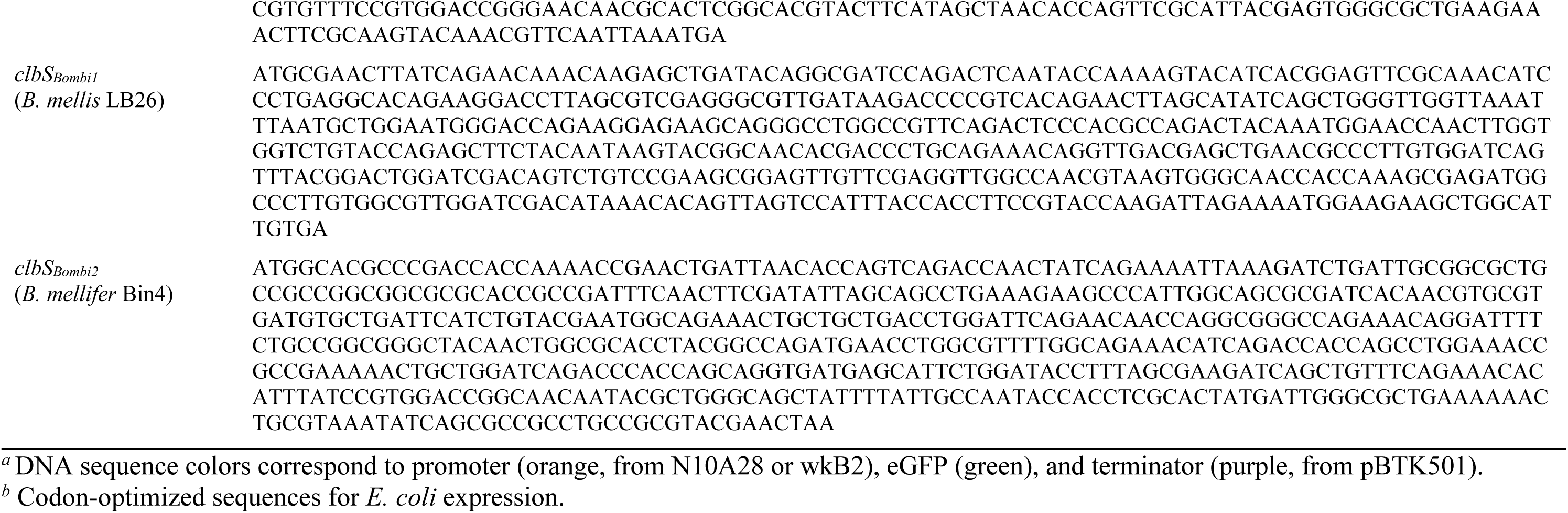
Synthesized DNA fragments. Gene block Sequence (5’ to 3’) Cassettes for DNA damage reporting*^a^*.

## Supplementary Data

**Data S1.** Presence of *clbS*-like genes in genomes of bee gut bacterial taxa and closely related taxa from other environments.

